# Neural voice activity detection with high-gamma ECoG signal correlation structure using a chronically implanted brain-computer interface in an individual with ALS

**DOI:** 10.1101/2025.08.21.671572

**Authors:** Kamil A. Grajski

## Abstract

Chronically implanted brain-computer interfaces (BCI) for speech have demonstrated restoration of speech communication for people with severe motor impairment. Yet maintaining stable long-term performance remains a translational challenge. We investigated whether correlation and partial correlation features of electrocorticographic (ECoG) high-gamma activity (HG-C, HG-PC) could improve robustness compared with high-gamma log-power (HGLP) features for neural voice activity detection (NVAD). Using open-source BCI data from an individual with amyotrophic lateral sclerosis performing a syllable-repetition task, long short-term memory (LSTM) models were trained separately on HGLP, HG-C, and HG-PC features, and evaluated across sessions spanning six months. HG-C and HG-PC achieved comparable or superior NVAD performance to HGLP with smaller long-term averaged loss (HGLP: -17%; HG-C: -9%; HG-PC: -12%) and smaller long-term worst case loss (HGLP: -24%; HG-C: -8%; HG-PC: -16%). Under a simulated local contiguous pattern of neural signal loss, both HG-C and HG-PC outperformed HGLP on averaged long-term loss (HGLP: -22%; HG-C and HG-PC, -10%) and worst-case long-term loss (HGLP: -30%; HG-C: -11%; HG-PC: -13%). The results show that high-gamma correlation-based features captured comparatively more spatially distributed and stable neural speech representations. With further refinement and validation, correlation-based feature representations may contribute to robust longitudinal speech decoding with implanted BCIs.

## Introduction

Implantable brain-computer interface (BCI) systems have enabled patients with paralysis to control external devices and to restore speech communication. Early implantable BCI demonstrations achieved reliable multidimensional cursor and robotic-arm control in tetraplegia [1–3], followed by rapid point-and-click communication and tablet interaction [4–6]. Recent clinical studies have extended these capabilities to speech decoding and speech synthesis in individuals with ALS, anarthria, dysarthria, or other severe speech impairments [7–15].

Implantable BCI device types vary by invasiveness and the recorded neural signal. Intracortical electrode grids such as the Utah grid [16, 17] and its variants are applied to record individual and small local neuron populations [9, 13, 15]. Sub-dural BCI devices embodied as linear strips [6] or grids [7, 10–12, 14] are applied to record cortical local field potentials as electrocorticograms (ECoG). An endovascular BCI is embodied in a multi-electrode cylindrical geometry [8, 18] for ECoG recording. A characteristic of chronically implanted BCI for ECoG recording is an inter-electrode center-to-center spacing in the range 1.0-4.0 mm, exposed electrode contact in the range 0.2-2.0mm, and electrode numbers in the single-digit to tens to 100-200. Higher-density electrode grids with sub-millimeter center-to-center spacing and contact area and electrode number up to 1024 have completed initial human intra-operative or temporary use in patients [19]. On a per device (e.g., multi-electrode grid) basis, these implantable BCI are designed to record neural signals from restricted cortical region(s). Principal implantation sites include the frontal lobe precentral gyrus primary motor cortex for device control and communication and ventral precentral gyrus for speech communication. Here, we focus on medium- to high-density implantable BCI for ECoG for speech communication.

Common to the operation of these BCI devices is a decoder. The decoder is a statistical model of the relationship between recorded neural signals and movement intention or speech production intention. Typical model inputs consist of windowed vector time-series of spike train-derived features or single- or multi-frequency band estimates of ECoG power. Model outputs take the form of a continuous control signal [1–5], discrete control signal [6], text (phoneme or syllable or word probabilities) [7, 9, 11, 45–48], text-to-speech [10, 13], or acoustic speech [12, 14, 15] when integrated into a complete system such as for at-home use. Decoder operation for at-home use typically entails an initial post-implantation training period comprised of multiple training sessions. Data collected during these sessions is stored for off-line neural signal processing and decoder training. Following training, model parameters initially are fixed and deployed to the BCI device for online real-time operations.

A key translational challenge for chronically implanted BCI is managing post-implantation model non-stationarity or “model drift”. Multiple factors contribute to model drift and each BCI device type is subject to one or more of these factors. Factors include neurodegeneration and atrophy of cortical motor areas such as that associated with ALS [34], long-term neural dynamics [20–26], and biological and mechanical effects [27–31]. While reliable performance can persist over years, it requires intermittent recalibration, adaptation, retraining, or self-calibration [3, 13, 23, 32–35, 36,37]. Robust feature engineering methods have been investigated for animal ECoG data [21–23] and recently applied to BCI [38–42].

Here, we propose a candidate robust decoder model derived from the ECoG high-gamma (HG) frequency band power signal associated with acoustic speech production intent [43–49]. The ECoG HG power signal undergirds recent studies in speech neuroprosthesis [7, 10–12, 14]. Specifically, we hypothesize that ECoG HG correlation and partial correlation structure is robust with respect to neural signal amplitude (power) drift and individual electrode and multi-electrode neural signal loss.

Two main considerations drive this hypothesis. First, BCIs based on processing multivariate EEG covariance with Riemannian geometric methods have been validated for feature representations, decoder design, and efficient calibration [50, 51]. These methods have been extended to functional connectivity estimators [52]. The typical target application for these approaches is for externally recorded whole-head EEG with inter-electrode spacing measured in centimeters. We find no prior EEG-Riemannian literature explicitly focused on application to the class of medium- to high-density ECoG BCI which are the focus of this study. Second, we observe that reported long-term changes in ECoG neural signal quality often appear to impact the amplitude of the high-gamma power envelope on individual or subsets of electrodes. We conjecture that, particularly for medium- to high-density electrode grids, local correlation structure - under the assumption that the neurophysiology itself remains functional long-term though possibly diminished - could still be estimated and exploited through correlation-based and partial-correlation features.

Here, we test the hypothesis by comparing short-term and long-term performance of a reference and a test neural voice activity detection (NVAD) model. The reference model is trained on HG (log) power envelope features (HGLP) and the test decoder is trained on HG-C (correlation-based) or HG-PC (partial correlation) features.

Here, we focused on neural voice activity detection as an initial and what we conjectured to be a straightforward binary classification task: assign the label of (non) speech frame by frame to windowed neural signal feature data. The study design was in four stages. The first stage was to reproduce as a baseline published findings obtained from the original open-source data [12]. This entailed training a neural network model comparable with that applied by Angrick, et al., using ECoG HGLP as input features and measuring performance (F1 score) as a function of elapsed time from last training data session.

The second stage was to generate and to explore ECoG correlation and partial correlation spatial and temporal structure. This stage answered basic questions. What does ECoG HG-C (HG-PC) data look like temporally and spatially? Could one visually observe the neural speech signal in ECoG HG-C (HG-PC)? Could one quantify the spatial extent of ECoG HG-C (HG-PC) such as through a spatial decay constant?

The third stage entailed training a neural network model of the same or similar architecture as in the first stage but utilizing HG-C (HG-PC). In this third stage, the ECoG HG-C (HG-PC) input feature data was first passed through a two-layer Linear Network for dimensionality reduction to match ECoG HGLP input dimensionality. Results at this stage include a comparison of HGLP, HG-C, and HG-PC NVAD performance stability and spatial distribution of marginal effects.

The final stage explored HGLP, HG-C, and HG-PC NVAD model performance in response to simulated neural signal loss.

## Methods

### Originating Study

A male, native English-speaking patient in his 60s with amyotrophic lateral sclerosis (ALS) was enrolled in a clinical trial (ClinicalTrials.gov **NCT03567213**) approved by the Johns Hopkins University Institutional Review Board (IRB) and conducted under an FDA Investigational Device Exemption. The trial evaluated the safety and preliminary efficacy of a BCI composed of subdural ECoG electrode arrays with a percutaneous connection to external amplifiers and computers. All procedures complied with relevant guidelines and regulations and were carried out according to the IRB-approved clinical protocol.

The neural data and anonymized speech audio data were obtained from the publicly available source (http://www.osf.io/49rt7) originated by Angrick, et al. (2024) [12]. The implantable BCI consisted of a pair of subdural 8x8 electrode grids (2.0 mm diameter; 4.0 mm center-to-center spacing). The device(s) were surgically implanted in the left hemisphere and placed on the pial surface of the sensorimotor representation areas of speech and upper extremities. Angrick, et al., applied this study data towards the estimation of models to generate online (real-time) speech synthesis from the 1000 Hz sampled electrocorticograms (ECoG). Their end-to-end system processed input comprised of digitally filtered ECoG high-gamma power (HGP) envelope activity. The system utilized three neural network stages: a) a unidirectional or causal neural network model that identified speech segments; b) a bidirectional model that translated the ECoG in voice-labelled segments to LPC coefficients; and c) LPCNet which converted LPC coefficients to acoustic speech.

### Study Data

The present study is an observational analytical secondary analysis of de-identified ECoG recordings available as open-source data by Angrick, et al. [12]. The original study and data were designed as a collection of thirteen (13) experimental sessions over a period of approximately seven months. We followed Angrick, et al., and organized the sessions into four groups. The training group consisted of the first eight sessions recorded over a one-month period. A single withheld independent testing session consisted of one session recorded approximately one week following the last training session. An additional single withheld independent validation session followed the testing session by one day. Finally, three sessions were recorded over the course of approximately one week starting approximately six months following the validation session. (See Supplementary Figure S1.)

Each recording session consisted of two sequential blocks of task-specific trials. The first block of trials consisted of a syllable repetition task (SRT). In an SRT trial, the participant was presented with an acoustic stimulus consisting of one of twelve consonant-syllable pairs. In random sequence, each of the twelve stimuli was presented five times, yielding a total of 60 trials per session. The second through n^th^ block of trials consisted of a keyword repetition task (KRT). In a KRT trial, the participant was prompted by a visual cue on a computer screen to read aloud one of six keywords or to remain silent. In random sequence, each of the six stimuli were presented ten times, yielding a total of 60 trials. Sessions contained at least one and up to eight KRT blocks.

In their study, Angrick, et al., utilized the SRT data for the purposes of gathering statistics to apply as baseline normalization of ECoG HGP features in the KRT data. Here, we focused exclusively on the SRT trials (N=780) as a source of NVAD training, testing, validation, and long-term stability testing. Metadata provided with the dataset included trial type, stimulus identifier, stimulus onset and offset times, and time-synchronized microphone audio recordings.

### Signal processing

Signal processing of ECoG data was implemented on a session-independent, block-independent, task-independent, trial-independent, and electrode channel-independent basis. There were three stages of signal processing. In the first stage, raw ECoG sample data time series were imputed for “bad channels” by simple nearest-neighbor spatial averaging. Angrick, et al., identified 0-3 bad channels which this study accepted. Additional detailed quality checking did not identify any additional bad channels.

In the second stage, pre-processing, the ECoG time series was centered to zero mean, digitally band-pass filtered to 70-170Hz, and then notch-filtered at 120Hz. The bandpass and notch filters were forward-backward digital filters using cascaded second order sections (SOS) which ensured that the output signals were temporally aligned with the input signals (including the audio signal). The output signal of the pre-processing stage was labelled the ECoG high gamma (HG) time series.

The third stage implemented feature extraction from the HG signal. First, the HG envelope magnitude was estimated using the Hilbert transform. Second, the windowed instantaneous high-gamma log power signal (HGLP) was estimated in a moving window (48 msec, duration; 12 msec, stride) by computing the mean of the squared magnitude and then taking log10. Each trial consisted of N=224 windows (frames). The selection of a 48 msec time window was based on the heuristic to include in each window at least 3-4 periods of signals at the lower end of the high-gamma frequency range. This study did not systematically consider other window and stride settings.

The resulting HGLP time series was then converted to Z-scores. Z-scoring ensured that all variables were on a similar scale with a mean of zero and a standard deviation of one, which prevents features with larger magnitudes from dominating the learning process and allows the neural network to converge faster during training. The resulting signal was referred to as HGLP feature data.

Windowed correlation-based features were extracted from the same HG time series used for HGLP estimation. For each window, the Pearson cross-correlation matrix was derived from the covariance matrix. The Pearson correlation coefficients were Fisher Z-transformed and stored as a strict lower triangular matrix. This signal was referred to as HG-C feature data.

Windowed partial correlation-based features were extracted from the same HG time series used for HGLP and HG-C estimation. For each window, for each electrode time series pair (**x**_i_, **x**_j_), their partial correlation was estimated by the Pearson cross-correlation of the residual time series pair **r**_i_, **r**_j_, where **r**_i_ was the residual of regression of **x**_i_ on all other electrode time series **x**_k_, except **x**_j_, and similarly for **r**_j_ . These correlation coefficients were also Fisher Z-transformed and stored as a strict lower triangular matrix. This signal was referred to as HG-PC feature data. HG-PC normalization was by quantile mapping to enhance the “burstiness” of the observed HG-PC signal by performing a two-stage transformation: robust baseline normalization followed by piecewise quantile-based contrast mapping. The robust baseline normalization computed mean and standard deviation using only the bottom 65% of amplitude values to prevent speech activity from contaminating the baseline statistics, while preserving the full variance for proper scaling. Piecewise quantile-based contrast mapping linearly mapped values within the 80th percentile to an output range of [0, 0.1], values between the 80th and 99th percentiles to [0.1, 1.0], and saturated values above the 99th percentile at the output value of 1.0. This approach enhanced burst contrast and maintained values in ranges suitable for neural network training. These values were determined as part of the neural network training hyperparameter grid search (see below).

When working with both 8x8 electrode grids, the dimensionality of the HGLP feature vector for each window was 128 and the dimensionality of the HG-C (HG-PC) feature vector was 8,128. When working with just one 8x8 electrode grid, such as the ventral grid, the HGLP feature dimension was 64 and the HG-C (HG-PC) dimension was 2,016.

Spatial correlation decay constants were estimated for HG-C and HG-PC (but not HGLP) by analyzing the relationship between inter-electrode distance and correlation estimate within each electrode grid (ventral and dorsal). Physical distances between same-grid electrode pairs were calculated using the known 8×8 grid geometry (4 mm inter-electrode spacing). For a given session and given stimulus the HG-C correlation values were pooled and grouped by exact the distance values (0, 4, 5.66, 8, 11.31… mm). An exponential decay model (ρ(d) = ρ₀ · ^(-d/λ)) was fitted to the distance-correlation data using least-squares optimization to extract the spatial decay constant λ.

### NVAD Labelling

This study leveraged the closed-loop nature of the SRT behavioral task to generate labelled training data for the NVAD task. Labelled training data consisted of assigning to each windowed feature vector (frame) the label 0 if “no speech” and 1 if “speech”. Voice Activity Detection (VAD) labeling was performed using microphone signal envelope analysis with the following methodology. For each trial, the microphone signal was processed using Hilbert envelope extraction and windowed to match ECoG signal processing. Reference statistics (mean and standard deviation) were computed from the pre-stimulus and stimulus periods (non-speech by SRT trial design). Speech frames were identified in the post-stimulus period using a threshold of k=1.5 standard deviations above the reference mean. Post-processing filled gaps between the first and last detected speech frames to enforce a single continuous speech segment. The decision to enforce one continuous speech segment per trial was based on the unitary nature of the consonant-syllable speech target. This policy potentially could introduce labeling errors as the patient displayed impaired articulation and phonation [12]. Following labelling, each trial was individually visually inspected and judged to be “correct” to within 1-2 window frames (12-24 msec) for speech onset and speech offset. To maximize reproducibility, there was no manual adjustment of frame labels. The labels were fixed once and only once at the start of the study and prior to any modeling or performance results from any feature representation. Summary statistics of the within-trial speech activity aggregated from the 780-trials in this study dataset are listed in Supplementary Table S1. No trials were excluded from the dataset. This approach achieved robust binary classification of speech vs. non-speech frames without requiring manual annotation. ECoG data was not referenced in any way during the automated labelling procedure.

### Neural Network Modeling

The neural network model selected was a stacked two-layer long short-term memory (LSTM) architecture with dropout between the layers followed by a linear fully connected output layer with two units (one signaling “speech”; the other signaling “no speech”). Ease of comparison with Angrick, et al., drove this decision. The nature of the NVAD task for SRT, including its time scale(s), dimensionality, and signal complexity, did not indicate more sophisticated architectures.

For HGLP models, the input layer was fixed at either 128 (dual grid) or 64 (single grid). For HG-C and HG-PC models, a multi-layer Linear Network (internal stages with ReLu output and dropout, but not final Linear Network stage) was incorporated ahead of the LSTM model. In the case of a single grid (ventral) model, the Linear Network mapped 2,016-dimensional input features to 64 output features then used as input to the LSTM model. In the case of a dual grid model, the Linear Network mapped 8,128-dimensional input features to 128 as input to the LSTM model.

Using Ray Tune with Optuna, an extensive hyperparameter search (Ray trial counts ranged 32-512 trials) was executed to estimate optimal signal pre-processing and network architectural and training parameters. For HGLP, HG-C, and HG-PC, the LSTM architectural hyperparameters included number of hidden units and dropout ratios. For HG-C and HG-PC, the Linear Network architectural hyperparameters included number of linear layers, number of hidden units per linear layer, and dropout ratio. For all feature types HGLP, HG-C, and HG-PC, training hyperparameters included batch size, number of training epochs, and learning rate (with fixed *ReduceLROnPlateau* parameters). We used Asynchronous Successive Halving Algorithm (ASHA) scheduler for early stopping during hyperparameter optimization. The ASHA scheduler was configured with an 8-epoch grace period, meaning trials were allowed to train for at least 8 epochs before being evaluated for early termination. Poorly performing trials were terminated early using a reduction factor of two, which significantly reduced computational costs during large searches. The scheduler monitored F1 score performance on the single first withheld test session only as the hyperparameter grid search optimization metric (see below). This allowed promising configurations to train to completion while terminating unpromising ones early.

The final ECoG HGLP LSTM model consisted of two layers with 256 hidden units each and 50%. dropout rate. The model was trained with batch size of 4, for 30 epochs, and learning rate 8.0 × 10⁻⁵. There were 856,578 trainable parameters.

The final ECoG HG-C LSTM model had the same architecture as for ECoG HGLP. The input Linear Reducer Network had 256 hidden units with 50% dropout. This mapped input 2,016 dimensions to 256 to 64 dimensions. The model was trained with batch size equal to 8, for 60 epochs, and learning rate 3.0 × 10⁻⁵. There were 1,389,378 trainable parameters total (856,578 (LSTM) + 532,800 (Reducer)).

The final ECoG HG-PC LSTM model consisted of two layers with 128 hidden units each and dropout 50% between layers. The input Linear Reducer Network had 512 hidden units with 50% dropout. This mapped 2,016 dimensions to 512 to 64 dimensions. The model was trained with batch size equal to 16, for 72 epochs, and learning rate 4.95 × 10⁻⁵. There were 1,297,218 trainable parameters total (231,424 (LSTM) + 1,065,536 (Reducer). HG-PC hyperparameter search assessed several alternative normalization methods and as noted above, quantile-mapping with the parameters reported above were incorporated into the final HG-PC model.

### Training and Performance Metrics

The neural network NVAD models were trained using only the eight training sessions. The training utilized cross-entropy loss estimation and the *RMSprop* optimizer implemented in PyTorch. LSTM internal state was reset to zero for each input trial. A standardized data representation permitted batch operations for rapid hyperparameter search, training, and inference.

The performance metric applied in this study was the F1 score for the per frame accuracy of classification of speech and no-speech. We noted that the study data had a class distribution of 75% non-speech and 25% speech frames. We judged this to be not severe enough to warrant additional processing such as to enforce an equal distribution. In addition to F1 score, AUC-PR was reported for each experiment.

Once each final model was trained with “champion” hyperparameters (no early stopping), the first single session of withheld independent test session data was used empirically to determine an optimal probability threshold with which to label a frame as “speech”. The probability threshold was selected as the value which maximized the test session F1-score. While, that probability typically defaults to *p*=0.5, we observed roughly 0.4 < *p* < 0.8. We did not undertake a systematic analysis of these values. Once computed, the probability threshold for each NVAD model type was frozen and applied in the evaluation of the succession of withheld sessions: independent validation and three online sessions. This strict-by-design method had the benefit of providing a basis to highlight inherent differential performance on sessions progressively removed in time from training sessions.

As a sanity check each of the models was retrained with fixed input features and hyperparameters, but with permuted frame labels in each trial. The F1-scores obtained on the test, validation, and online sessions were in the range 0.39 - 0.44 roughly corresponding to the geometric mean of the class distribution, as expected.

As a final check, each of the models was retrained twenty (20) times with fixed input features and hyperparameters, but different initial random seed. The main effect of different random number seeds was on the composition of samples into batches, the sequencing of input data, and the initialization of weights and drop-out.

### Marginal EAects

To assess the performance of the final trained NVAD models, we computed individual feature level marginal effects. The marginal effect of an individual feature was defined as the change in test set F1 score when that individual feature was randomly permuted while all others were held constant: baseline test set F1 score minus permuted test set F1 score. Feature sample permutation preserved marginal distribution while its relationship to signal was randomized. A positive value indicated that permuting the feature degraded performance, implying it was “useful” to the model. The distribution of test set F1 scores under model training with correct labels was compared to that of the same model trained with permuted target labels (see above). The combination of a model trained with permuted labels and then analyzed by permuting individual features confirmed the null distribution of marginal effects as zero.

### Simulated Signal Loss

To simulate performance impact of post-training neural signal loss, we replaced recorded neural signals with “noise” in feature space. In the case of HGLP, neural signal loss was simulated by replacing individual or multiple HGLP feature dimension(s) with random samples drawn from the Normal distribution *N*(0,σ=1) for all trials in all sessions evaluated by an NVAD final model. In the case of HG-C (HG-PC), neural signal loss on an electrode was simulated by replacing each of the (partial) correlation pairs involving that electrode with *N*(0,σ=1) random samples for all trials in all session evaluated. For clarity, the final NVAD models themselves were not perturbed and the sessions used for evaluation were the test session, the validation session, and the three “online” sessions. (See Supplementary Figure S1.) This study explored three patterns of neural signal loss: single electrode; multi-electrode local contiguous grid; and randomly distributed multi-electrode signal loss.

In the localized neural signal loss approach, individual electrodes or small local neighborhoods (1×1, 3×3, 5×5 grids) of electrodes were replaced with N(0,σ=1) noise while keeping the remaining signals intact. This operation was repeated such that every electrode in the grid served as a center electrode position. Per session, per grid size results were summarized as the mean and standard deviation of F1 score difference from baseline performance with intact signals aggregated across the electrodes in the grid tested (e.g., ventral).

In the random neural signal loss approach, the mean and standard deviation of session F1 scores were computed over repeated runs where the number of runs was at least 10, but up to 200 and adaptively determined based on the number of possible random subsets of a given size in the range tested (N=1-16). Performance degradation was measured as percentage change in F1 score from fully intact baseline. These analyses were conducted for HGLP, HG-C, and HG-PC models. As a control, the procedure was also performed on NVAD models trained with permuted target data.

### Statistical Analysis

Statistical analysis was selectively conducted to evaluate session effects or feature effects on NVAD F1-score baseline and simulated neural signal loss performance differences. For baseline analysis, session differences within each feature type were assessed using repeated-measures ANOVA with Mauchly’s test for sphericity and Greenhouse–Geisser correction when violated. For feature effects, differences were evaluated per session using one-way ANOVA when assumptions were met, or Kruskal-Wallis H-test when normality was violated. Statistical significance was defined at α = 0.05 for omnibus tests, with Bonferroni correction applied to post-hoc pairwise comparisons to control family-wise error rate.

Post-hoc tests used paired *t*-tests (repeated measures), independent *t*-tests (one-way ANOVA), Wilcoxon signed-rank tests (when normality assumptions were violated for repeated measures), or Mann-Whitney U tests (Kruskal-Wallis). Effect sizes for all pairwise tests were quantified using Cohen’s *d*, interpreted according to standard conventions (|*d*| < 0.2 negligible, 0.2 ≤ |*d*| < 0.5 small, 0.5 ≤ |*d*| < 0.8 medium, |*d*| ≥ 0.8 large). All analyses were implemented in Python using scipy.stats and statsmodels packages.

## Results

We defined a standardized trial and focused on NVAD of the acoustic speech recorded during the Syllable Repetition Task (SRT). Figure 1 shows a representative trial with HGLP feature data. We trained an HGLP NVAD model with N=128 input features (dorsal and ventral electrode grids) and performed a marginal effects analysis. (See Supplementary Figure S2.) We found marginal effects to be largely, but not completely comparable to the findings reported by Angrick, et al., who used a partial derivative-based saliency method [53]. Where Angrick, et al., registered meaningful saliency for four electrodes in the dorsal grid (Figure 3 in [12]), our method measured progressively more negative marginal effect with distance from the ventral region of the ventral grid. We noted that the partial derivative method ignores inter-feature dependencies, may saturate in nonlinear regions, depends on local data distribution, and varies with feature scaling. Further, such open-source information that was available did not reflect the dorsal and ventral grid placement, orientation, and distance details necessary independently to assess the neurophysiological plausibility of incorporating data from both electrode grids into a single model. While we explored HGLP, HG-C, and HG-PC data from dorsal and ventral electrode grids, here we report and statistically analyze NVAD model performance results based on neural signal recordings from the ventral 8x8 electrode grid only.

**Figure 1.**
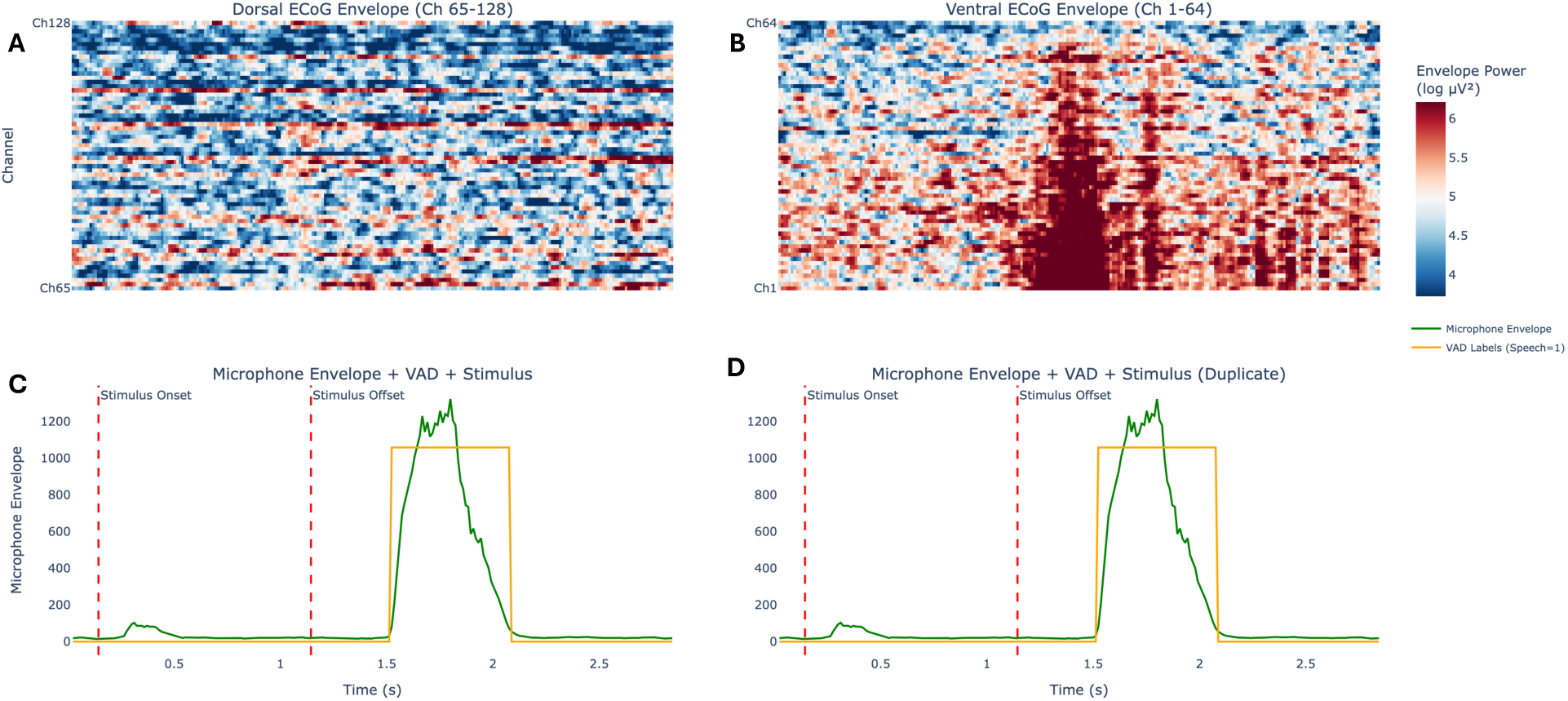
Standardized syllable repetition task neural voice activity detection trial example (*BAH*). A standardized trial spans the time 144 msec pre-stimulus, continuing through the 1.0 second stimulus delivery period, and ending 1728 msec post-stimulus. In each of thirteen (13) sessions, the patient completes a syllable repetition task comprised of randomized trials (N=60) 2.5-3.5 seconds duration resulting in five (5) repetitions of twelve (12) consonant-vowel audio stimuli played through a loudspeaker. **(A)** HGLP from dorsal 8x8 electrode grid. **(B)** HGLP from 8x8 electrode grid. **(C)** and **(D)** Identical for ease of comparison with HGLP plots above. Stimulus onset and offset (*dotted red lines*); microphone sample data windowed average (time-aligned with HGLP) (*solid green line*); and automatically labelled speech ON/OFF (0/1) target label (*orange line*). A key observation is the visual evidence of neural signal correlates of voice activity.

Table 1 summarizes results for NVAD model performance based on HGLP, HG-C, or HG-PC. Performance was measured as F1 score (geometric mean of the accuracy of correctly labelling individual trial windows (frames) as speech or non-speech) and as area-under-curve precision-recall (AUC-PR) per session. Results are listed for each of the sessions in the sequence of post-training sessions as mean and standard deviation over repeated runs (N=20) of the final trained models using unique random number seeds.

**Table 1.** Performance comparison for HGLP, HG-C, and HG-PC NVAD models. The Session column lists the session identifying label together with number of weeks elapsed from last training session shown in parentheses. **(A)** F1 Score. **(B)** AUC-PR. Columns show the mean and standard deviation aggregated over twenty repeated runs of each of the respective final models with unique random number seeds per run. The results are based on NVAD model output alone. No post-processing or filtering or padding or smoothing of predicted labels was applied. This is by design to isolate the contributions of the neural signal representation on performance. Supplementary Table S2 lists details of the statistical analysis of these results.

| **A** | Session | HGLP F1 Score | HG-C F1 Score | HG-PC F1 Score |
| --- | --- | --- | --- | --- |
|  | Test (1.0) | 0.87 ± 0.006 | 0.89 ± 0.013 | 0.89 ± 0.022 |
|  | Validation (1.1) | 0.85 ± 0.006 | 0.89 ± 0.017 | 0.87 ± 0.021 |
|  | Online 1 (24.1) | 0.77 ± 0.009 | 0.82 ± 0.016 | 0.80 ± 0.031 |
|  | Online 2 (24.7) | 0.74 ± 0.013 | 0.79 ± 0.019 | 0.79 ± 0.031 |
|  | Online 3 (25.1) | 0.66 ± 0.018 | 0.82 ± 0.016 | 0.75 ± 0.037 |
| **B** | Session | HGLP AUC-PR | HG-C AUC-PR | HG-PC AUC-PR |
|  | Test (1.0) | 0.94 ± 0.005 | 0.96 ± 0.006 | 0.97 ± 0.009 |
|  | Validation (1.1) | 0.93 ± 0.007 | 0.96 ± 0.005 | 0.95 ± 0.017 |
|  | Online 1 (24.1) | 0.83 ± 0.008 | 0.93 ± 0.008 | 0.88 ± 0.033 |
|  | Online 2 (24.7) | 0.79 ± 0.013 | 0.90 ± 0.017 | 0.87 ± 0.034 |
|  | Online 3 (25.1) | 0.72 ± 0.024 | 0.89 ± 0.012 | 0.82 ± 0.048 |

The F1 score performance percentage change for HGLP, HG-C, and HG-PC was -24%, -8%, and -16%, respectively, measured as last online session (online 3, approximately +25 weeks from the last training session) compared with the test session (approximately +1 week from the last training session). The F1 score performance percentage change for HGLP, HG-C, and HG-PC was -17%, -9%, and -12%, respectively, measured as the average of the last three online sessions (online 1, 2, and 3) compared to the test session. Statistical analysis explored session effects and feature effects. Session effects were highly significant for all three feature types (all *p* < 0.001), with HGLP showing the largest effect size (*F* = 1089.9), followed by HG-C (*F* = 137.7) and HG-PC (*F* = 83.0). Post-hoc comparisons demonstrated significant performance differences between most session pairs, with particularly large effect sizes for Test vs Online comparisons (Cohen’s *d* > 10 for HGLP). Feature effects analysis revealed significant differences across feature types in all sessions (all *p* < 0.001). HG-C outperformed HGLP across all sessions, with effect sizes ranging from moderate (*d* = -1.68, test) to very large (*d* = -8.96, online 3). HG-PC showed intermediate performance, generally outperforming HGLP but underperforming HG-C in later online sessions. (See Supplementary Table S2.)

HGLP, HG-C, and HG-PC qualitatively demonstrated characteristic error patterns. HGLP-based NVAD gave rise to false alarms that resulted in one or more predicted speech sequences within a single trial or a single predicted sequence completely offset from ground-truth. In contrast, HG-C and HG-PC-based NVAD characteristic errors primarily related to timing of onset and offset of the ground-truth speech segment. (See Supplementary Figure S10.)

The remainder of this section presents an analysis of the feature data, marginal effects, and model robustness in response to neural signal loss.

### NVAD with ECoG Power Features (HGLP)

The marginal effects procedure was applied to the ventral electrode grid HGLP NVAD model. Although 2/3 of HGLP features (electrodes) had positive marginal effects, there was an order of magnitude difference in marginal effects value between the top-ranked and 10^th^ ranked feature.

This reflected a concentration of marginal effects to a small subset of available features. (See Supplementary Figure S3.)

### NVAD with ECoG Power Correlation Features (HG-C)

The standard trial provided a convenient framework with which to perform initial data visualization of ECoG HG-C feature data. Figure 2 shows overlayed HG-C time series plots for the ventral grid. Also shown is the spatially sorted HG-C data, including both dorsal and ventral grids. (See Supplementary Figure S4 for dorsal grid and dorsal ventral inter-grid HG-C time series overlays.)

**Figure 2.**
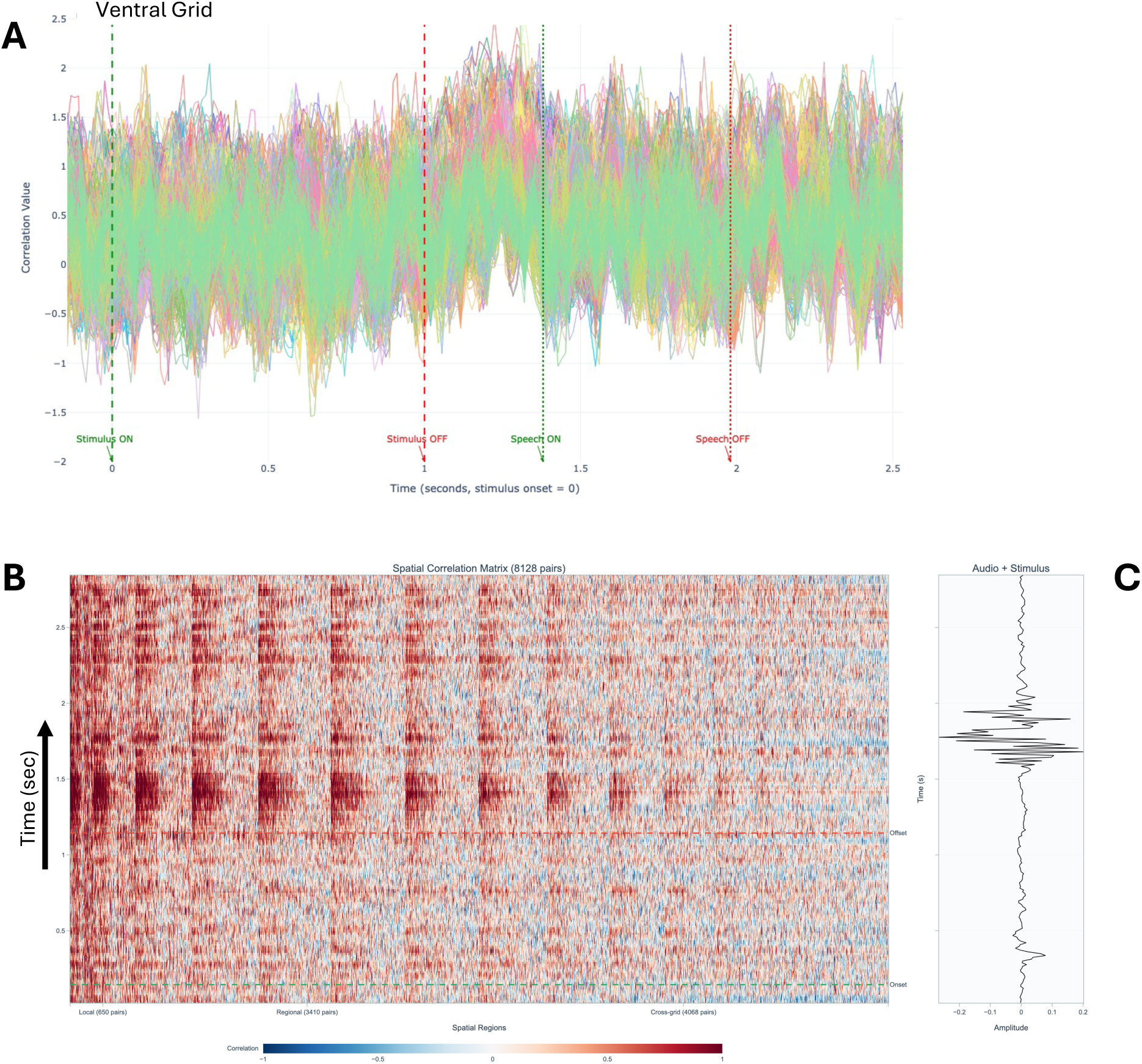
HG-C in a standardized neural voice activity detection trial (*same as* Figure 1 *trial*). **(A)** HG-C overlaid time series plots for the ventral intra-grid electrode pairs. The *x*-axis shows time from trial start in seconds. The *y*-axis shows Fisher Z-transformed Pearson correlation coefficient of windowed HGLP time series. A key observation is the increase in correlation values immediately post-stimulus and preceding speech onset. The corresponding plots for dorsal intra-grid and cross-grid ventral and dorsal are shown in Supplementary Figure S4. **(B)** Spatial-distance sorted HG-C structure including HG-C from both ventral and dorsal grids. The y-axis shows time from trial start (*bottom left*) to 1,768 msec post stimulus (*top left*). The x-axis shows individual Pearson correlation coefficient pairs of the HGLP windowed time series. Dotted horizontal green and red lines (at right of panel) mark stimulus onset and offset. **(C)** Windowed audio average signal (*x-axis*) time-aligned (*y-axis*) with spatial correlation structure shown at left. A key observation is the spatial structure evident at both local (near-neighbors, at left of plot panel (A)), regional (distant within-grid, middle portion of plot panel (A)), and to a visually far lesser extent between grid distance scales (right portion of plot panel (A)) immediately preceding, during, and following the patient’s consonant-vowel voicing.

A quantitative spatial analysis was performed to characterize the visually observed HG-C spatial signature with an estimated spatial decay constant measured in millimeters (mm). Figure 3 shows typical results for one stimulus (“BAH”) across the five individual SRT trial repetitions and averaged response recorded during a single training session. The ventral spatial decay constant (38-48mm) was 4-5x longer than the dorsal spatial decay constant (8-10mm), irrespective of pre-stimulus, stimulus, and post-stimulus time periods. The ventral post-stimulus period showed an increase in correlation strength and longer spatial decay compared to stimulus and pre-stimulus periods.

**Figure 3.**
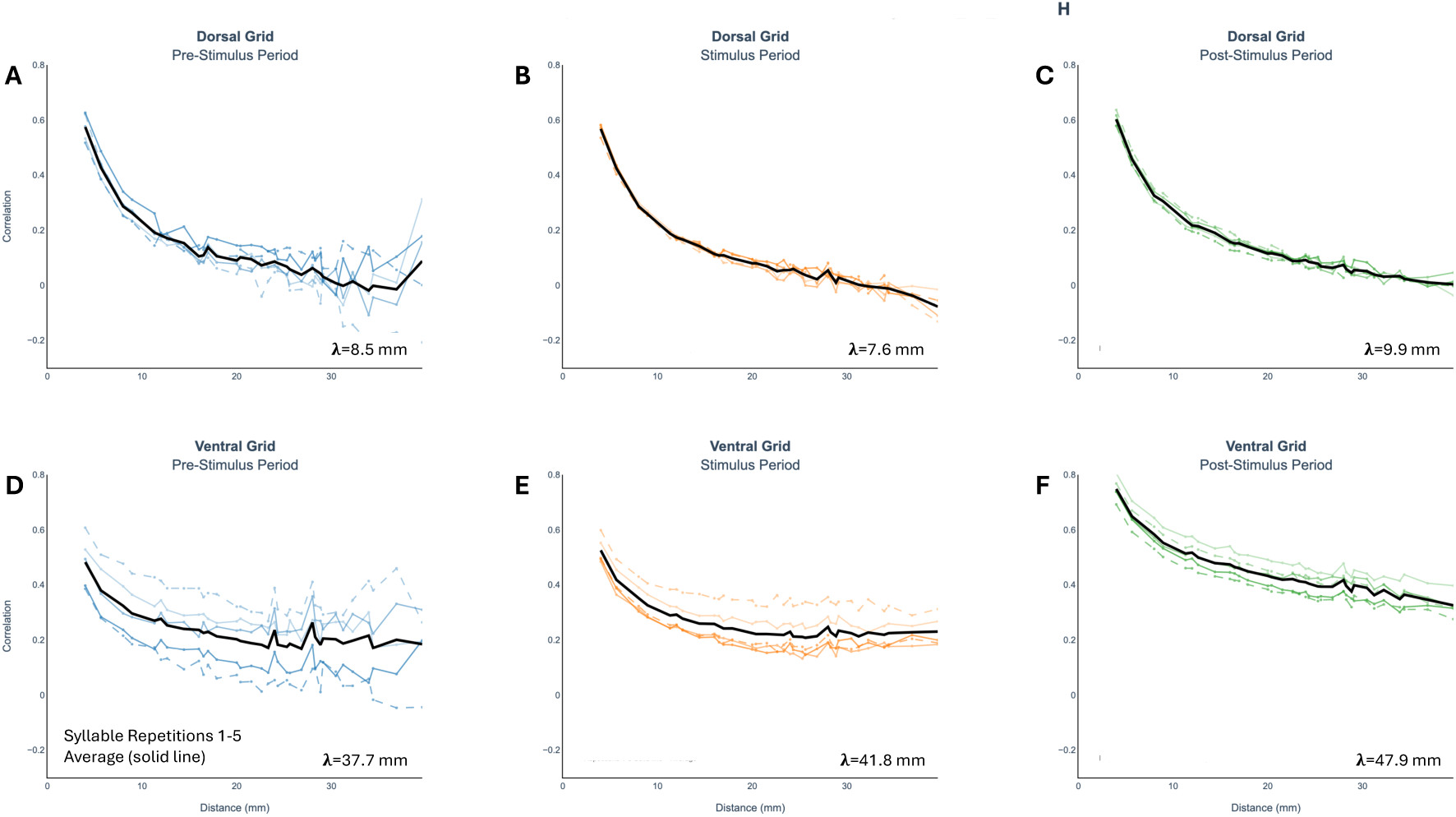
Standardized neural voice activity detection trial time periods showing empirically estimated HG-C spatial decay constants. Each panel shows results for each of the five utterances of the given stimulus (BAH) recorded during one session. The *y*-axis shows Pearson correlation coefficient values. The *x*-axis shows distance in units of mm between electrode pairs. **(A)** Dorsal grid pre-stimulus period. **(B)** Dorsal grid stimulus period. **(C)** Dorsal grid post-stimulus period. **(D)** Ventral grid pre-stimulus period. **(E)** Ventral grid stimulus period. **(F)** Ventral grid post-stimulus period. The dorsal and ventral grid analyses are conducted independently as they are physically embodied as a pair of 8x8 electrode grids offset in distance and rotated one with respect to the other. The results shown are typical across stimulus types, trials, and sessions.

The marginal effects procedure was applied to the ventral electrode grid HG-C NVAD model. The spatial distribution of HG-C marginal effects is shown in Figure 4. (See Supplementary Figure S5 for the marginal effect distribution and ranked distribution plots.) HG-C marginal effects were comparable in part with findings with HGLP. Correlation pairs that included HGLP high marginal effect values also had elevated marginal effect values (e.g., electrode 21 Supplementary Figure S3). However, HG-C feature marginal effects were more widely distributed. This is shown visually by plotting individual feature marginal effect values relative to the global average marginal effect. The marginal effects procedure was applied also to the HG-C NVAD model trained with permuted target labels - thus permuting both feature data and labels - with the expected zero marginal effects per feature (not shown).

**Figure 4.**
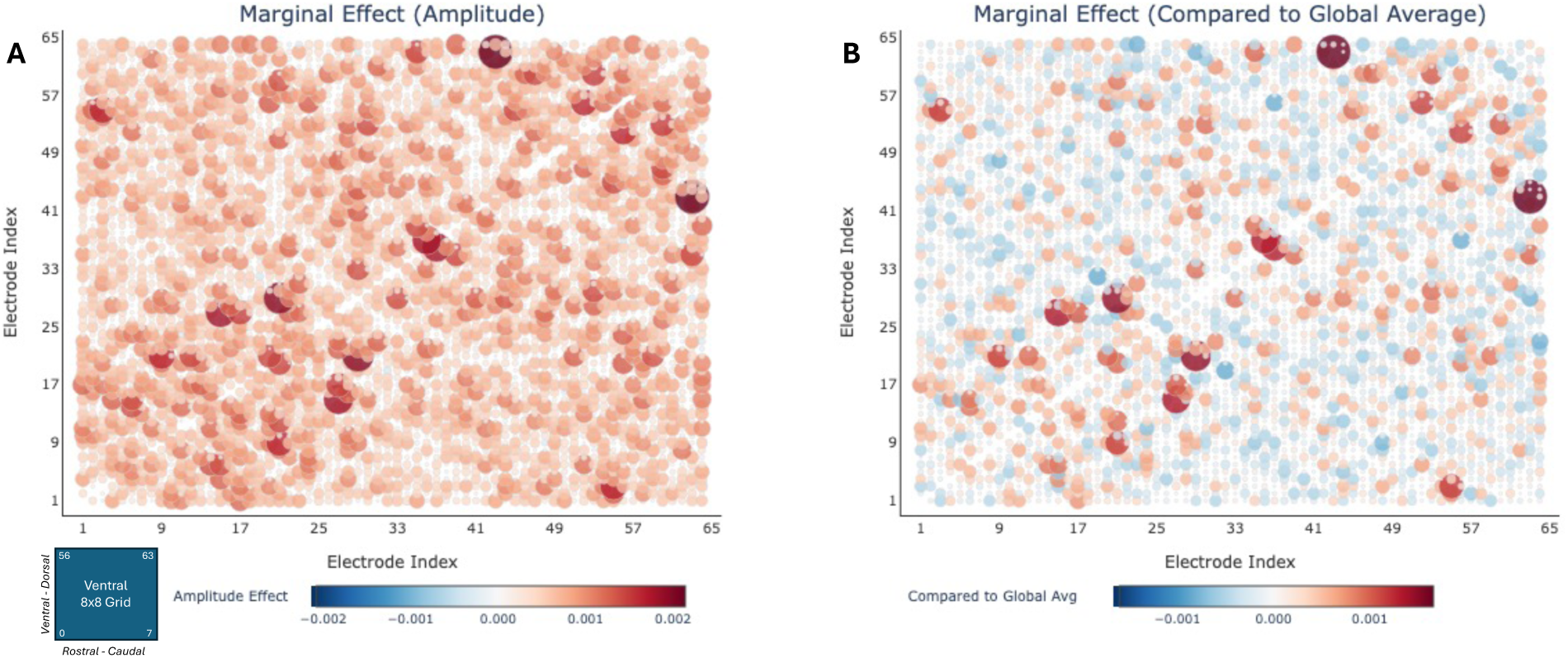
Spatial distribution of empirical marginal effect of individual HG-C features on test session F1 score performance of the HG-C NVAD model. Each subplot is in the form of a symmetric correlation matrix. Each point in the subplot represents a pairwise correlation computed in the ventral 8x8 electrode grid in a single typical trial. The correlation matrix is rotated to maintain the convention in the Angrick, et al., study data that electrode #1 is in the rostral- and ventral-most position in the ventral electrode grid. **(A)** Pair-wise correlation marginal effects when NVAD model is trained with correct target labels. **(B)** The same pair-wise correlation marginal effects plotted as values relative to the global average correlation marginal effect. Diameter and color proportional to relative marginal effect amplitude. A key visual observation is the multiplicity of high marginal effect locations.

### NVAD with ECoG Power Partial Correlation (HG-PC)

The results of the HG-C spatial decay analysis suggested that there could be shared variance in the signal(s) in the ventral and dorsal HG-C. See Figure 3. As a first step, we undertook a partial correlation analysis to remove shared variance explained by all other ventral and dorsal electrodes. The standard trial provided a convenient framework with which to perform initial data visualization of ECoG HG-PC raw data. Figure 5 shows overlays of the 2,016 HG-PC time series plots for the partial correlation pairs in the ventral 8x8 electrode grid. Also shown is the spatial structure spanning both ventral and dorsal electrode grids. The “burstiness” of the HG-PC time series signal is more pronounced than either HGLP or HG-C. Unexpectedly, visual inspection of the HG-PC time-series overlays and spatial structure revealed comparatively stronger activity within the dorsal grid and with inter-grid connections than was observed for HGLP or HG-C. (See Supplementary Figures S4 and S6.) This pattern may indicate additional distributed coupling sampled within and between the dorsal and ventral electrode grids. The HG-PC spatial decay curves had lower average value and were “flat” as expected (*not shown*). In the second step of HG-PC analysis in this study, we re-estimated HG-PC, but strictly with and within the ventral electrode grid. The results reported in Table 1 are on such basis. The marginal effects procedure was performed on the HG-PC and we observed comparable findings of wider distribution of marginal effects compared to HGLP (See Supplementary Figure S7).

**Figure 5.**
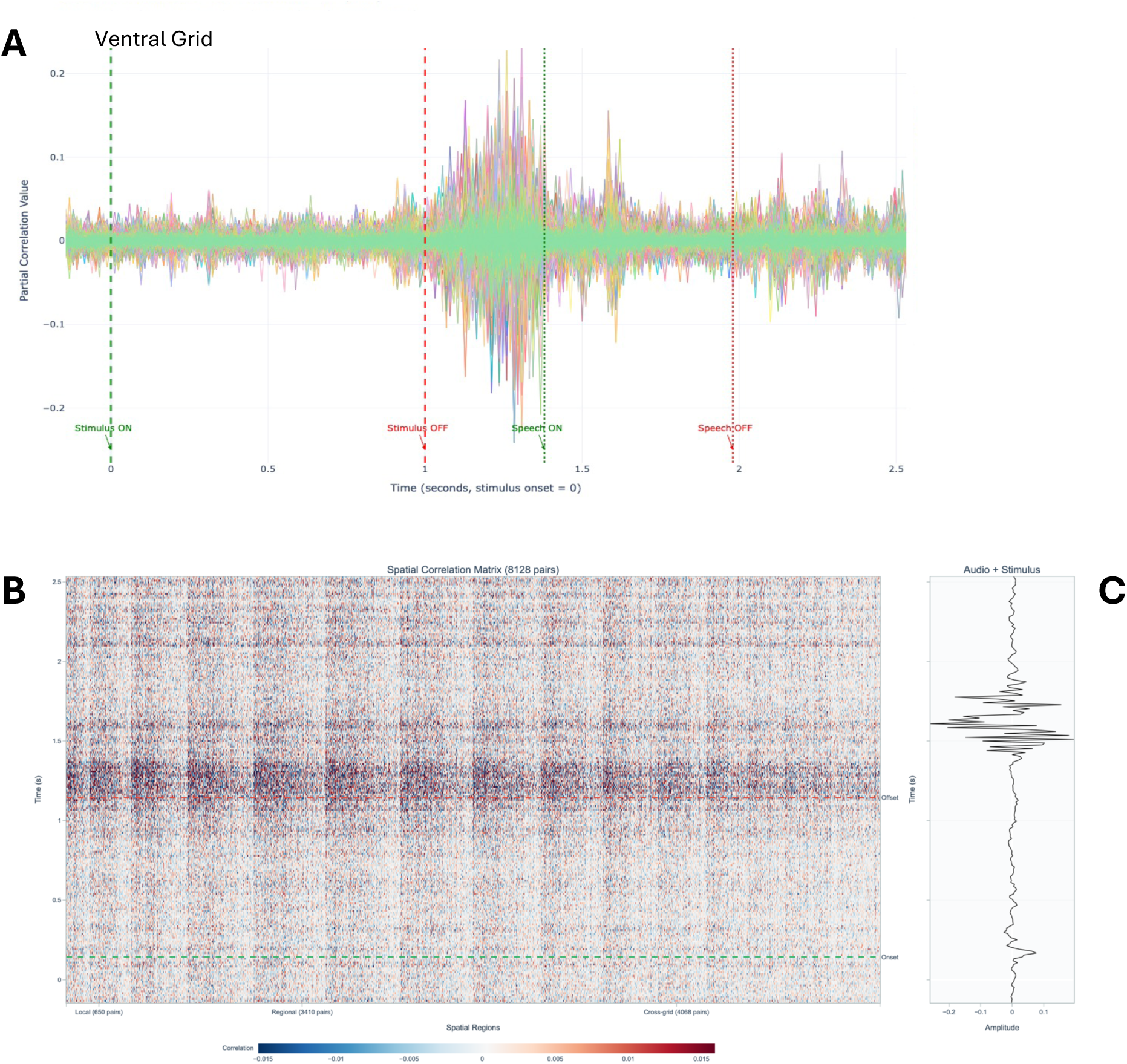
HG-PC in a standardized neural voice activity detection trial (*same as* Figure 1 *trial*). **(A)** HG-PC overlaid time series plots for the ventral intra-grid electrode pairs. The *x*-axis shows time from trial start in seconds. The *y*-axis shows Fisher Z-transformed partial Pearson correlation coefficient of windowed HGLP time series. A key observation is the “burst” in correlation values immediately post-stimulus and preceding speech onset. The corresponding plots for dorsal intra-grid and cross-grid ventral and dorsal are shown in Supplementary Figure S6. **(B)** Spatial-distance sorted HG-C structure including HG-PC from both ventral and dorsal grids. The y-axis shows time from trial start (*bottom left*) to 1,768 msec post stimulus (*top left*). The x-axis shows individual partial Pearson correlation coefficient pairs of the HGLP windowed time series. Dotted horizontal green and red lines (at right of panel) mark stimulus onset and offset. **(C)** Windowed audio average signal (*x-axis*) time-aligned (*y-axis*) with spatial correlation structure shown at left. A key observation is the spatial structure evident at both local (near-neighbors, at left of plot panel (A)), regional (distant within-grid, middle portion of plot panel (A)), and to a visually far greater extent compared to HGLP and HG-C at the between grid distance scales (right portion of plot panel (A)) immediately preceding, during, and following the patient’s consonant-vowel voicing..

### Simulated Neural Signal Loss (HGLP, HG-C, HG-PC)

To test NVAD model robustness, we simulated neural signal loss in individual and locally contiguous subsets of electrodes. Summary results are shown in Figure 6A-6C. For the 3x3 multi-electrode neural signal loss experiment (9 electrodes at a time out of 64 set to *N*(0,1) noise) measured over the last three “online” sessions, HG-C and HG-PC NVAD outperformed HGLP with smaller long-term averaged loss (HGLP: -22%; HG-C: -10%; HG-PC: -10%) and smaller long-term worst case loss (HGLP: -30%; HG-C: -11%; HG-PC: -13%).

**Figure 6.**
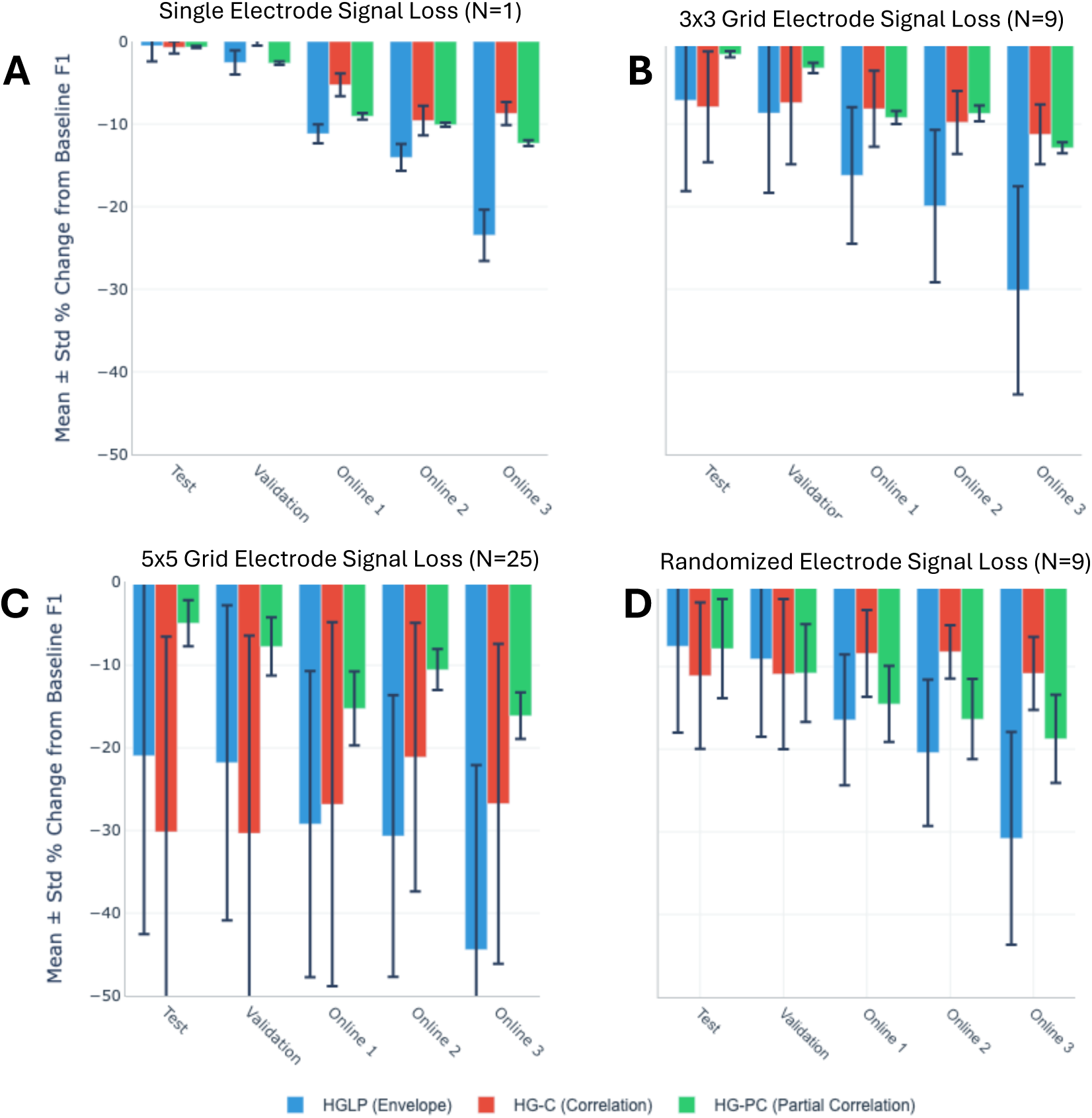
Effect of simulated neural signal loss on NVAD F1score. The plots show in summary the results of four simulated neural signal loss experiments. (A) Single electrode. (B) 3x3 contiguous grid. (C) 5x5 contiguous grid. (D) Random spatial distributed set of nine electrodes - to serve as comparison with the 3x3 grid case shown in (B). For (A), (B), and (C), the plots show for each session the mean and standard deviation percentage change in F1 score compared to intact model aggregated across the 64 center electrodes. For (D), the plot shows for each session the mean and standard deviation percentage change in F1 score compared to intact model aggregated across repeated (N=200) random samples of size nine (9).

Statistical analysis of feature effects in independent local contiguous grid experiments revealed significant effects across all grid sizes tested and sessions (all omnibus tests *p* < 0.001). For the 1×1 grid, HGLP consistently underperformed both HG-C and HG-PC across all sessions, with very large effect sizes (Cohen’s *d* ranging from -1.87 to -7.18). The 3×3 grid showed similar patterns but with reduced effect magnitudes, while the 5×5 grid demonstrated more variable significance, particularly for HGLP vs HG-C comparisons in early sessions. (See Supplementary Table S3).

We hypothesized that for a model that learns primarily local features (HGLP) and under local contiguous neural signal loss, the variance of performance would be higher compared to models that learn distributed features (HG-C, HG-PC). For the 3x3 neural signal loss case, the standard deviation (read as percentage points of degradation) averaged over the three long-term sessions was: HGLP: 10.0% HG-C: 4.0%; HG-PC: 0.8%. That is, HGLP variation in performance with position of neural signal loss was 2.5 times greater than that of HG-C and 12.5 times greater than that of HG-PC. (See Supplementary Figure S8.)

To test whether robustness to neural signal loss extended beyond spatially contiguous outages, we performed randomized neural signal loss experiments. (See Supplementary Figure S9). We observed degraded performance in all feature types. We conducted detailed statistical analysis of feature effects in several independent random electrode count experiments (1, 3, 5, 9). We observed in all cases, significant effects in the omnibus tests (*p* < 0.01) for online sessions. Post hoc paired tests were significant exclusively for the HGLP, HG-C pair, but with small effect (typically for online 1 and online 2) to medium effect (typically for online 3) as measured by Cohen’s *d*. We compared the 3x3 grid loss case (Figure 6B) to the equivalent number of electrodes randomly selected (N=9) (Figure 6D). For HG-PC, but not HGLP and not HG-C, the per session variance of performance degradation in the grid case was a fraction of the performance degradation in the random case.

## Discussion

### Summary

We hypothesized that correlation features of electrocorticographic high-gamma activity could improve robustness compared with high-gamma log-power features. By design, this study focused on the well-studied and relatively straightforward task of neural voice activity detection. These initial results - based on data from a single patient performing a single task over an approximately seven month period and with model parameters fixed post-training - show that ECoG HG-C and HG-PC features support neural voice activity detection with performance comparable to and in key respects superior to HGLP features. HG-C and HG-PC models generated significantly less performance degradation over time. HG-C held greater resilience to both local contiguous and random patterns of neural signal loss compared to HGLP and HG-PC. These findings show that correlation-based high-gamma features captured spatially distributed long-term stable representations. With further refinement and validation, such representations could potentially contribute to robust longitudinal speech decoding with implanted BCIs.

### Limitations

There are limitations in this study. The task itself, consonant-vowel repetition, may be too restrictive, artificial, and of limited clinical value. A related limitation is that the dataset is small, a closed-loop design, and idiosyncratic to one patient. As a result, there is risk of overtraining and of over-interpretation of errors. The methods applied in this study sought to manage overtraining risk through systematic hyperparameter tuning, aggressive dropout rates in the LSTM model, and in the case of HG-C and HG-PC input, the Linear Model stage as well. Over-interpretation of errors is a risk as the types of errors generated by the NVAD models could be managed with threshold adjustments and post-processing of frame labels. Finally, the comparison between HGLP and HG-C (HG-PC) did not control for number of trainable parameters. During hyperparameter grid search, however, far higher trainable parameter counts were evaluated and we report here results with the optimal architectures, data batching, and training parameters.

### Refinement and Validation

Refinement and validation of the methods and results reported here include consideration of additional NVAD model architectures, exploration of additional ECoG frequency bands, application of Riemannian methods, validation with additional larger ECoG datasets, and potentially the extension to other neural signal data modalities.

The HG-PC results reported here show that distributed correlation structure may extend beyond the ventral grid to include dorsal regions. This indicates development and evaluation of NVAD architectures that utilize correlation-based methods across both electrode grids.

This study focused on ECoG high-gamma frequency band by design as it is a well-understood reference against which any improvements must be measured. One possibility is to operate with both alpha and gamma bands such as adopted by Vansteensel, et al., [6, 34]. Synergistic with multi-band analysis is the opportunity systematically to explore window size and stride impacts other than those applied here.

As noted, Riemannian methods have been successfully applied in EEG BCI [50–52]. Application here requires redesign of the workflow including of normalization, model training, marginal effects analysis, and simulated neural signal loss in a fashion that preserves Riemannian geometry throughout. For further ECoG studies, there is additional data available from the Angrick, et al., study analyzed here that is comprised of a closed loop keyword repetition task (KRT) with six command words [12].

Extension of the methods reported here to additional neural signal modalities with open-source data could include stereotactic EEG (sEEG) and intracortical microelectrode neural signal data. The confirmed available open-source Verwoert, et al., dataset is comprised of closed-loop sEEG recordings using 1,103 electrodes obtained in ten patients as part of their clinical treatment for epilepsy and who spoke 100 Dutch language words and numbers [54]. Finally, the confirmed available open-source Willet, et al., dataset is comprised of intracortical microelectrode recordings from an ALS patient who had lost his ability to speak and demonstrated high-performance with a speech neuroprosthesis [9, 55].

## Code Availability

All source code supporting this study will be made publicly available on https://github.com/kgrajski/ecog_stability upon acceptance of the manuscript.

## Data Availability

The raw time series samples of neural data and anonymized speech audio were previously made publicly available at http://www.osf.io/49rt7/ by the original study team [12]. The present study produced: data refactored as standardized trials, feature data comprised of ECoG HGLP and correlation matrix-based features derived from HGLP (HG-C, HG-PC) as individual trials stored as .csv files; a metadata database for task, session, block, and trial metadata; and a database of experimental results. These may be made available upon reasonable request to foster collaboration.

## Author Contributions

KAG conceived of and designed the experiments to assess ECoG HG-C and HG-PC reported in this manuscript, implemented all code underlying the processing methods and results, analyzed the data, and wrote the manuscript.

## Acknowledgements

Acknowledgements first and foremost to the ALS patient whose enrollment in the clinical trial (NCT03567213) resulted in the creation of this study dataset. Acknowledgements and appreciation to the Nathan E. Crone Lab and Miguel Angrick at the Department of Neurology at the Johns Hopkins University School of Medicine who elected to make even the raw time series study dataset available as open source.

