## Supplementary Material for "Neural voice activity detection with high-gamma ECoG signal correlation structure using a chronically implanted brain-computer interface in an individual with ALS"

| NVAD Standardized Syllable Repetition Task (SRT) Per Trial Speech Segment Statistics From Automated Labelling | | | |
|---|---|---|---|
| Cumulative Statistics | Trials With Acoustic Speech (N, %) | Acoustic Speech Latency (s) | Acoustic Speech Duration (s) |
| All Trials (N=780) | 777, 99% | | |
| Mean, Std. Dev. | | 0.967, 0.248 | 0.913, 0.229 |
| Range | | 0.108 - 2.412 | 0.048 - 1.668 |

**Table S1.** NVAD standardized Syllable Repetition Task trial automated acoustic speech labelling statistics. The participant was instructed to repeat the audio prompt immediately upon termination of an audio prompt. The study dataset consisted of 60 recorded trials from each of 13 sessions (N=780 trials). An automated voice activity labelling method, followed by visual inspection, was used to create "ground truth" speech activity. Ninety-nine percent of trials contained speech. There were three trials with no speech and five trials with speech cutoff at end of trial. These trials remained in the study dataset. At the trial frame level (224 per trial; 174,720 total) approximately 25% of frames were labelled as speech and 75% as non-speech. This was judged to be moderately imbalanced, but not so imbalanced as to require special handling (e.g., up-sampling, down-sampling, case weights, class weights, etc.).

**A**

| Comparison | HGLP | HG-C | HG-PC |
|---|---|---|---|
| Omnibus Test (*F, p*) | *F=1089.9, p<0.001* | *F=137.7, p<0.001* | *F=83.0, p<0.001* |
| Test vs Validation (*t, p, d*) | *t=12.19, p<0.001, d=2.73* | - | *t=6.29, p<0.001, d=1.41* |
| Test vs Online 1 (*t, p, d*) | *t=57.50, p<0.001, d=12.86* | *t=17.00, p<0.001, d=3.80* | *t=16.25, p<0.001, d=3.63* |
| Test vs Online 2 (*t, p, d*) | *W=0.00, p<0.001, d=10.68* | *t=16.49, p<0.001, d=3.69* | *t=18.90, p<0.001, d=4.23* |
| Test vs Online 3 (*t, p, d*) | *t=51.74, p<0.001, d=11.57* | *t=15.88, p<0.001, d=3.55* | *t=25.93, p<0.001, d=5.80* |
| Validation vs Online 1 (*t, p, d*) | *t=41.68, p<0.001, d=9.32* | *t=18.39, p<0.001, d=4.11* | *t=19.30, p<0.001, d=4.31* |
| Validation vs Online 2 (*t, p, d*) | *t=45.00, p<0.001, d=10.06* | *t=15.97, p<0.001, d=3.57* | *t=19.47, p<0.001, d=4.35* |
| Validation vs Online 3 (*t, p, d*) | *t=51.70, p<0.001, d=11.56* | *t=13.82, p<0.001, d=3.09* | *t=25.54, p<0.001, d=5.71* |
| Online 1 vs Online 2 (*t, p, d*) | *t=12.59, p<0.001, d=2.82* | *t=8.08, p<0.001, d=1.81* | - |
| Online 1 vs Online 3 (*t, p, d*) | *t=30.03, p<0.001, d=6.71* | - | *t=11.96, p<0.001, d=2.67* |
| Online 2 vs Online 3 (*t, p, d*) | *W=0.00, p<0.001, d=5.15* | *t=-6.38, p<0.001, d=-1.43* | *t=10.23, p<0.001, d=2.29* |

**B**

| Session | Omnibus Test | HGLP vs HG-C | HGLP vs HG-PC | HG-C vs HG-PC |
|---|---|---|---|---|
| Test | $\chi^2$*=17.5, p<0.001* | *W=10.00, p<0.001, d=-1.68* | - | - |
| Validation | $\chi^2$*=22.8, p<0.001* | *W=2.00, p<0.001, d=-2.76* | - | - |
| Online 1 | $\chi^2$*=22.8, p<0.001* | *W=0.00, p<0.001, d=-3.87* | - | - |
| Online 2 | $\chi^2$*=24.4, p<0.001* | *W=0.00, p<0.001, d=-3.38* | *W=3.00, p<0.001, d=-2.41* | - |
| Online 3 | *F=175.0, p<0.001* | *t=-28.35, p<0.001, d=-8.96* | *t=-9.21, p<0.001, d=-2.91* | *t=7.24, p<0.001, d=2.29* |

**Table S2.** Statistical analysis of NVAD F1-score performance across sessions and feature types. **(A)** Session effects. Within-feature comparisons across sessions (read by column). **(B)** Feature effects. Within-session comparisons across feature types (read by row). All *p*-values are Bonferroni-corrected for multiple comparisons. Results are based on twenty repeated runs of the respective final models with unique random number seeds per run. All omnibus tests were significant; empty cells indicate non-significant pairwise tests ($p > 0.001$). Note: *F* = *F*-statistic from repeated measures ANOVA; *t* = *t*-statistic from paired *t*-tests; *W* = Wilcoxon signed-rank test statistic; *p* = *p*-value (Bonferroni-corrected); *d* = Cohen's *d* effect size.

| Grid Size | Session | F-statistic | HGLP vs HG-C | HGLP vs HG-PC | HG-C vs HG-PC |
|---|---|---|---|---|---|
| 1x1 | Test | *H=164.86, p<<0.001* | *U=118.00, p=0.000, d=-1.87* | *U=0.00, p=0.000, d=-2.91* | *U=4.00, p=0.000, d=-2.19* |
| | Validation | *H=164.64, p<<0.001* | *U=0.00, p=0.000, d=-4.83* | *U=0.00, p=0.000, d=-3.71* | *U=3968.00, p=0.000, d=3.05* |
| | Online 1 | *H=161.30, p<<0.001* | *U=0.00, p=0.000, d=-6.71* | *U=0.00, p=0.000, d=-6.72* | *U=3880.00, p=0.000, d=2.74* |
| | Online 2 | *H=131.69, p<<0.001* | *U=0.00, p=0.000, d=-4.10* | *U=0.00, p=0.000, d=-6.56* | - |
| | Online 3 | *H=161.56, p<<0.001* | *U=0.00, p=0.000, d=-7.18* | *U=0.00, p=0.000, d=-6.66* | *U=3887.00, p=0.000, d=2.53* |
| 3x3 | Test | *H=126.45, p<<0.001* | - | *U=0.00, p=0.000, d=-1.23* | *U=15.00, p=0.000, d=-1.58* |
| | Validation | *H=95.40, p<<0.001* | *U=1208.00, p=0.000, d=-0.46* | *U=0.00, p=0.000, d=-1.38* | *U=871.00, p=0.000, d=-1.00* |
| | Online 1 | *H=109.32, p<<0.001* | *U=304.00, p=0.000, d=-1.59* | *U=0.00, p=0.000, d=-1.82* | - |
| | Online 2 | *H=123.51, p<<0.001* | *U=130.00, p=0.000, d=-1.80* | *U=0.00, p=0.000, d=-2.28* | - |
| | Online 3 | *H=127.78, p<<0.001* | *U=19.00, p=0.000, d=-2.31* | *U=0.00, p=0.000, d=-2.33* | - |
| 5x5 | Test | *H=86.67, p<<0.001* | - | *U=619.00, p=0.000, d=-1.29* | *U=186.00, p=0.000, d=-1.57* |
| | Validation | *H=69.11, p<<0.001* | - | *U=597.00, p=0.000, d=-1.30* | *U=503.00, p=0.000, d=-1.38* |
| | Online 1 | *H=41.05, p<<0.001* | - | *U=649.00, p=0.000, d=-1.29* | - |
| | Online 2 | *H=105.63, p<<0.001* | *U=992.00, p=0.000, d=-0.70* | *U=10.00, p=0.000, d=-1.95* | *U=734.00, p=0.000, d=-1.00* |
| | Online 3 | *H=97.42, p<<0.001* | *U=853.00, p=0.000, d=-0.94* | *U=0.00, p=0.000, d=-1.99* | *U=1156.00, p=0.000, d=-0.84* |

**Table S3.** Statistical analysis of F1-score performance differences among HGLP, HG-C, and HG-PC feature types across five experimental sessions and three systematic grid sizes (1x1, 3x3, 5x5). Each grid size was analyzed independently. The repeated measures for F1-score performance changes for a given session, feature, and grid size were aggregated across 64 experiments in which each electrode in the ventral 8x8 electrode grid was the "center" electrode in the loss grid. All omnibus tests were significant; empty cells indicate non-significant pairwise tests ($p > 0.001$). Note: $F$ = one-way ANOVA $F$-statistic; $H$ = Kruskal-Wallis $H$-statistic; $t$ = independent t-statistic; $U$ = Mann-Whitney $U$-statistic; $d$ = Cohen's $d$ effect size; $p$ = p-value (Bonferroni-corrected).

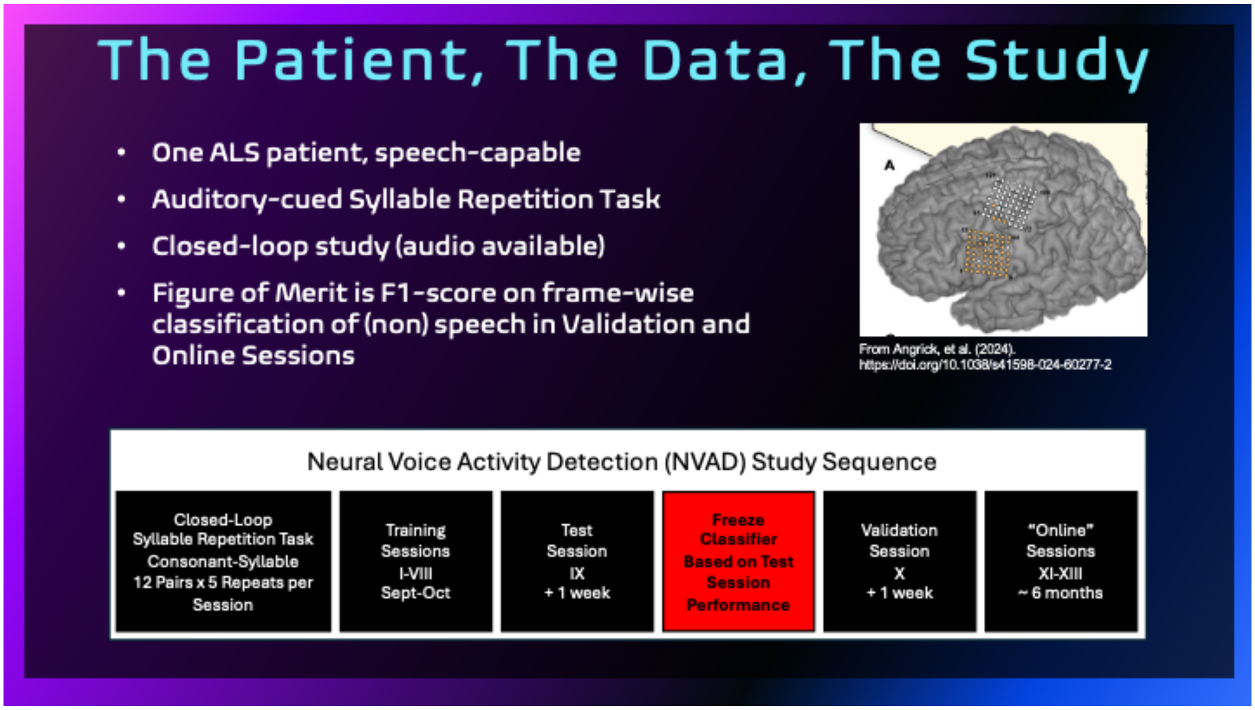

**Figure S1.** Overview of the patient, the data, and the study. Inset (*top right*) is from Angrick, et al., showing approximate placement of the pair of 8x8 electrode grids referenced as the ventral and dorsal electrode grids. The bottom row shows chronological progression and elapsed times of Syllable Repetition Task sessions recorded with the patient. Eight sessions were recorded over the period of a few weeks in October and November and used as training data. An independent withheld test session was recorded approximately one week following the last training data session as was an additional independent validation session. The three "online" sessions followed by approximately six months. F1 score performance on the test session was optimized by adjusting the probability threshold used to classify frames as speech. Thereafter all parameters of the model were fixed. By fixing all model parameters following test session, the workflow enables estimation of immediate (validation session) and medium- to long-term (online sessions) performance.

### Marginal Effects for Model Trained with BOTH Dorsal and Ventral Electrode Grids
### Distribution, Ranked Feature Importance, and Spatial Distribution

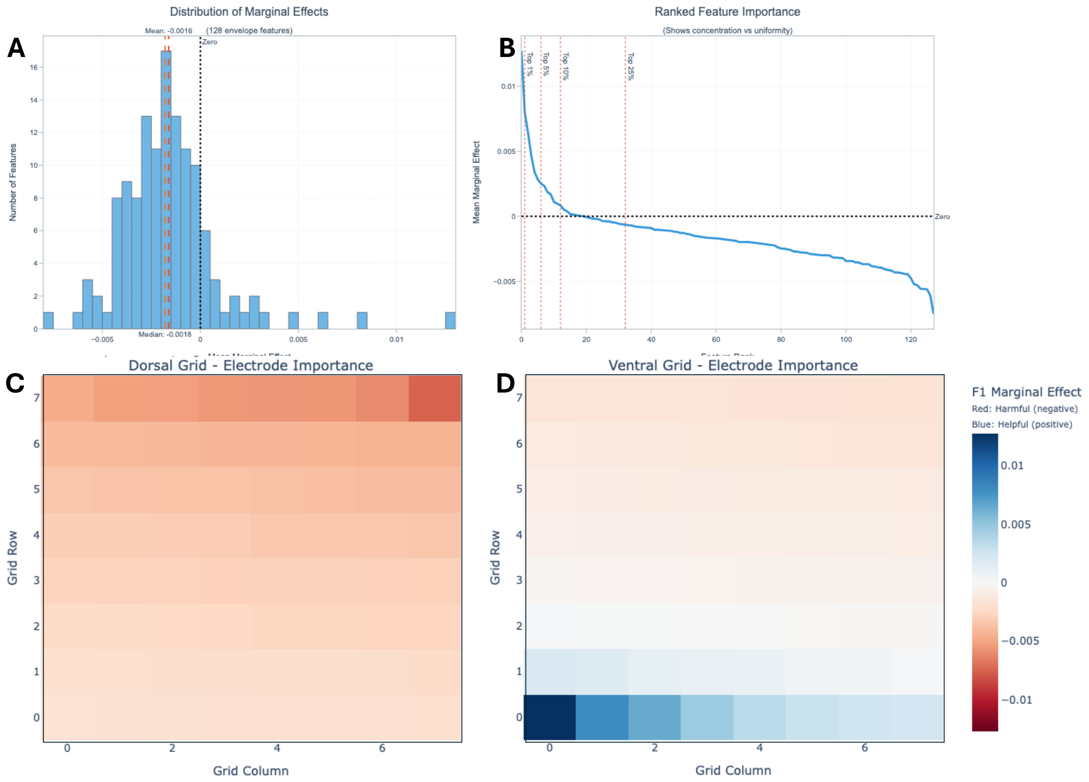

**Figure S2.** Distribution, ranked distribution, and spatial distribution of individual electrode HGLP empirical marginal effects on test session F1 score performance of an HGLP NVAD model trained with dorsal and ventral neural signals. **(A)** Histogram of values; mean and median score (*dotted lines*). **(B)** Ranked values; percentiles (*dotted red lines*). A key observation is that the distribution of empirical marginal effect scores has a long positive tail. As a control, the NVAD model training procedure was repeated with target labels permuted and then followed by the marginal effects procedure - thus permuting labels and feature values - which results were the constant zero for individual electrode marginal effect as expected. **(C)** Spatial distribution of marginal effects of values for dorsal 8x8 grid. **(D)** Spatial distribution of marginal effects of values for ventral 8x8 grid. We observe in this data that all positive-valued marginal effects are concentrated in the ventral-most region of the ventral grid. As distance grows from the ventral-most portions of the grid and proceeding dorsally, the marginal effects smoothly decay towards zero and then turn increasingly negative. Consequently, in this study, model training and evaluation proceeded using only ventral neural data.

### Marginal Effects for Model Trained with ONLY Ventral Electrode Grid Distribution, Ranked Feature Importance, and Spatial Distribution

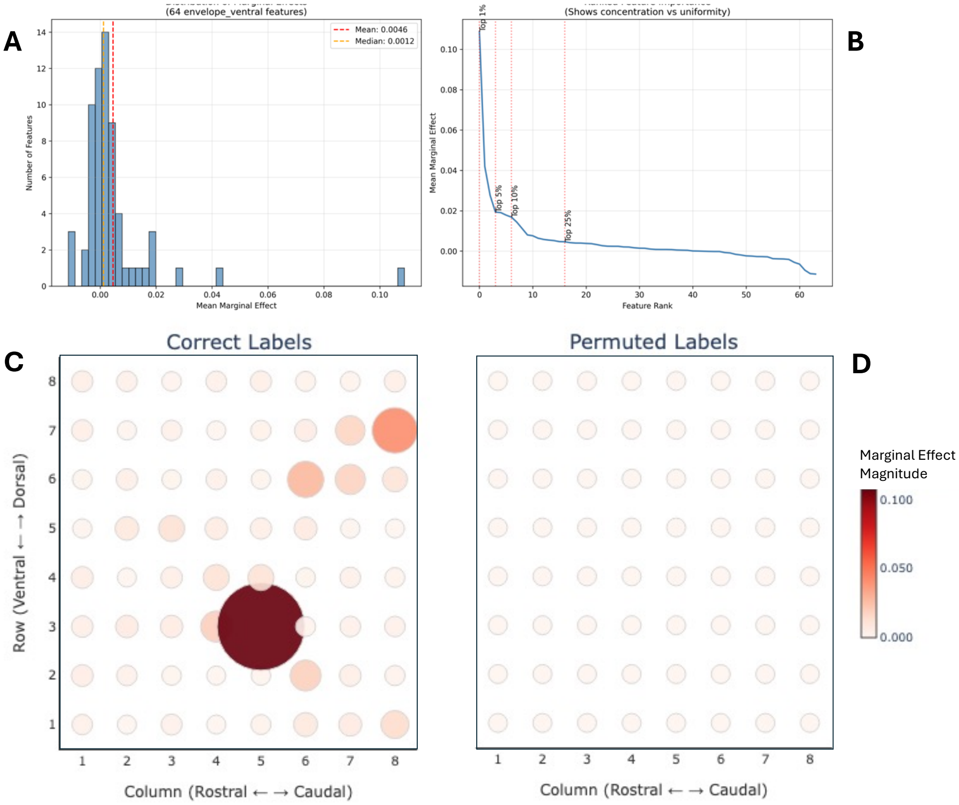

**Figure S3.** Distribution, ranked distribution, and spatial distribution of individual electrode HGLP empirical marginal effects on test session F1 score performance with the ventral grid HGLP NVAD model. **(A)** Histogram of values; mean and median score (*dotted lines*). **(B)** Ranked values; percentiles (*dotted red lines*). A key observation is that the distribution of empirical marginal effect scores is non-uniform with the top 5$^{th}$-10$^{th}$ percentile of "excessively" important electrodes (relatively high positive value). **(C)** Marginal effects heatmap of values for ventral 8x8 electrode grid when HGLP NVAD model is trained with correct target labels. **(D)** Marginal effects heatmap of values for ventral 8x8 electrode grid when HGLP NVAD model was trained with permuted target labels - thus permuting both features and labels - confirmed expected result of zero marginal effects. Note: Circle diameter and color proportional to marginal effect magnitude.

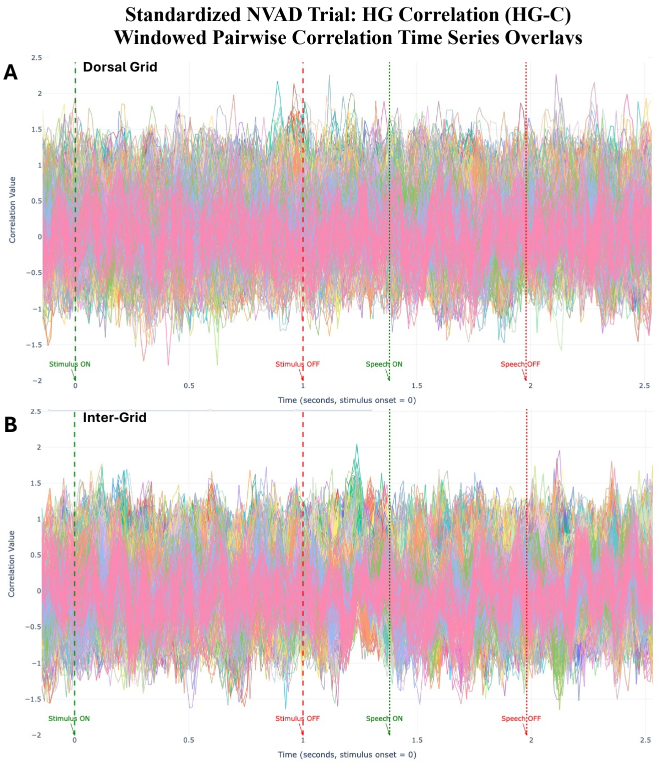

**Figure S4.** Time series visualization of HG-C in a standardized neural voice activity detection trial (*same as Figure 1 trial*) shows overlays of individual windowed pairwise correlation time series plots. **(A)** Dorsal intra-grid electrode pairs. **(B)** Inter-grid electrode pairs. The *x*-axis shows time from trial start in seconds. The *y*-axis shows Fisher Z-transformed Pearson correlation coefficient of windowed HGLP time series. Visually, there appears to be less change from pre-stimulus baseline in the peri-stimulus and acoustic speech time periods compared to the ventral grid (Figure 2).

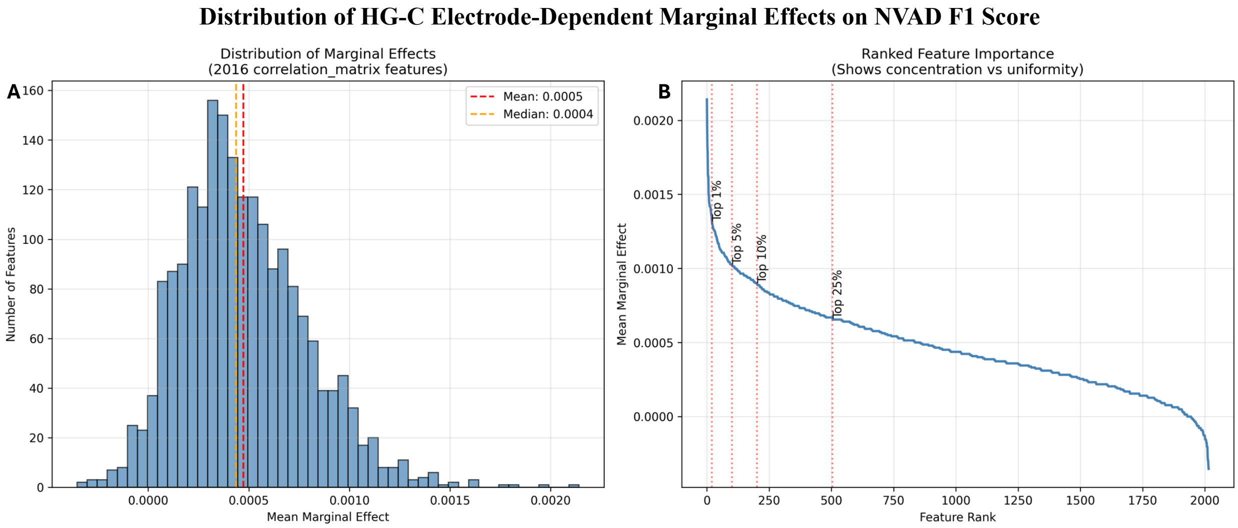

**Figure S5**. Distribution and ranked distribution of empirical marginal effect values of individual ventral 8x8 electrode grid pairwise correlation HG-C features (N=2,016) on test session F1 score performance of the NVAD model. **(A)** Histogram of values; mean and median score (*dotted red lines*). **(B)** Ranked values; percentiles (*dotted red lines*). A key observation is that the distribution of empirical marginal effect scores is non-uniform with a top $5^{th}$-$10^{th}$ percentile of very "important" electrodes (relatively high positive value). The NVAD model training procedure was repeated with permuted target labels and then subjected to the marginal effects procedure – thus permuting both labels and feature data - which yielded the expected constant zero for individual HG-C feature (correlation pair) marginal effect.

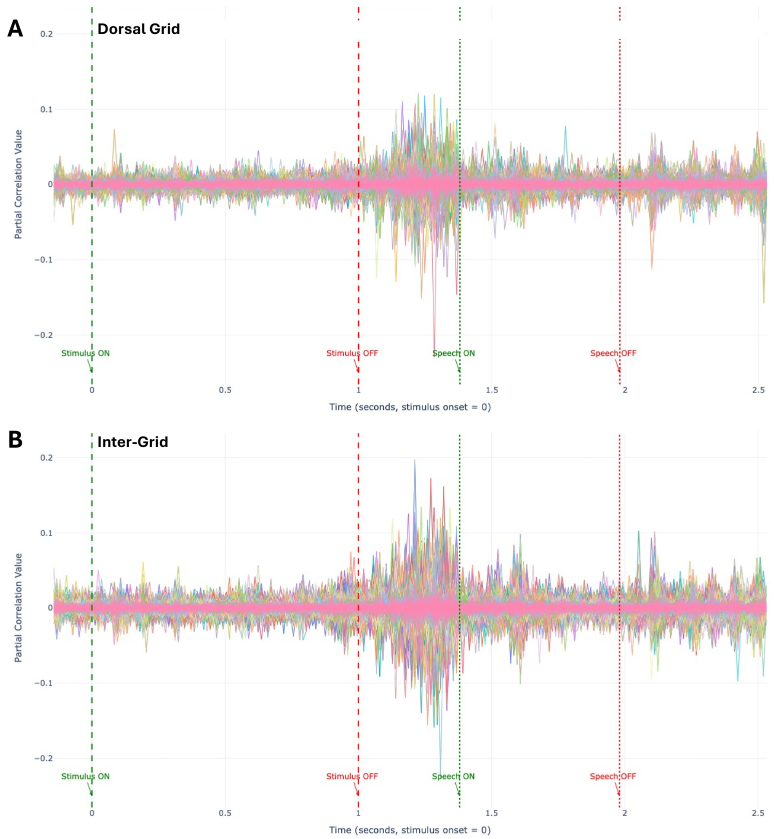

**Figure S6.** HG-PC in a standardized neural voice activity detection trial (*same as Figure 1 and Figure 3 trial*) shows overlays of individual windowed pairwise partial correlation time series. **(A)** Dorsal intra-grid electrode pairs. **(B)** Inter-grid electrode pairs. The *x*-axis shows time from trial start in seconds. The *y*-axis shows Fisher Z-transformed Pearson correlation partial coefficient of windowed HGLP time series. A key observation is the increase in correlation values immediately post-stimulus and preceding speech onset. Contrary to what was visually observed with HGLP and HG-C, there is visually apparent HG-PC activity in the period post-stimulus to pre-acoustic speech in both the dorsal and inter-grid plots. This observation was not further pursued in this study.

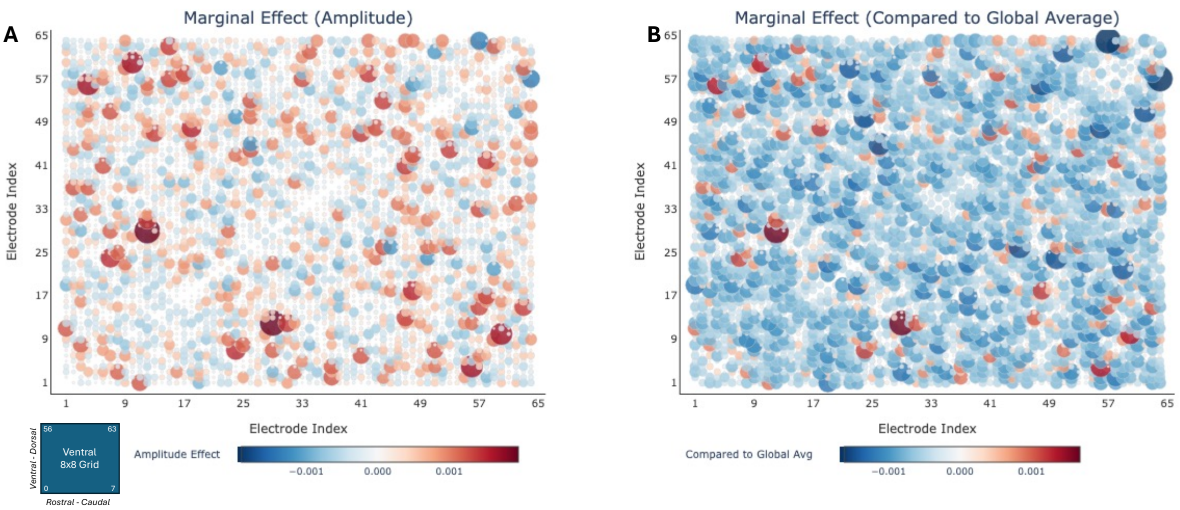

**Figure S7**. Spatial distribution of empirical marginal effect of individual HG-PC features on test session F1 score performance of the HG-PC NVAD model. Each subplot is in the form of a symmetric correlation matrix. Each point in the subplot represents a pairwise partial correlation computed in the ventral 8x8 electrode grid in a single typical trial. The correlation matrix is rotated to maintain the convention in the Angrick, et al., study data that electrode #1 is in the rostral- and ventral-most position in the ventral electrode grid. **(A)** Pair-wise correlation marginal effects when NVAD model is trained with correct target labels. **(B)** The same pair-wise partial correlation marginal effects plotted as values relative to the global average correlation marginal effect. Diameter and color proportional to relative marginal effect amplitude. A key visual observation is the multiplicity of high-magnitude marginal effect locations.

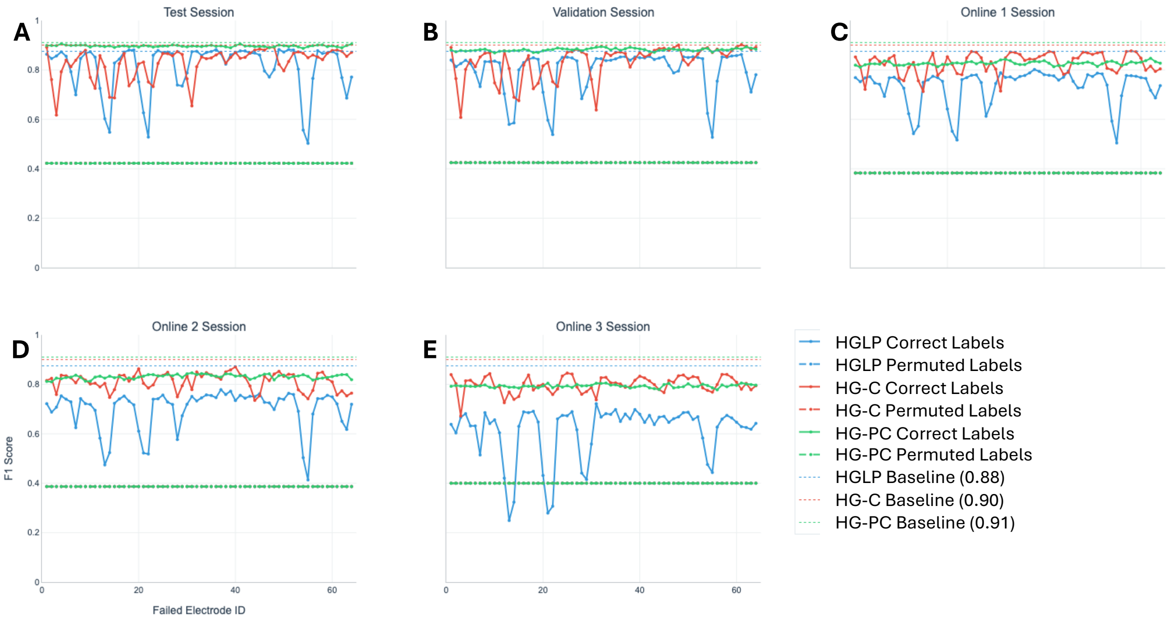

**Figure S8.** Details of the effect of simulated neural signal loss on NVAD F1 score by session for successive contiguous local 3x3 grid signal loss in the ventral grid. Neural signal loss is simulated by systematically setting all HGLP values or all pairwise HG-C (HG-PC) values to $N(0, \sigma=1)$ in the N x N grid (N=1, 3, 5) centered at the $i^{th}$ electrode in the range 1 to 64 (ventral grid only) and evaluating NVAD performance. Along electrode grid edges, the simulated failure grid size is less than N x N. For each panel in the Figure – one panel per session - we show: HGLP NVAD trained with correct labels (blue solid line) and an HGLP NVAD model trained with permuted target labels (blue dashed horizontal line at ~0.4); HG-C NVAD trained with correct labels (red solid line) and an HG-C NVAD model trained with permuted target labels (red dashed horizontal line at ~0.4); HG-PC NVAD trained with correct labels (green solid line) and an HG-PC NVAD model trained with permuted target labels (green dashed horizontal line at ~0.4); and the F1 score obtained on Test set by the HGLP, HG-C, and HG-PC NVAD intact models (shown as dotted blue, red, and green lines near the top of each subplot, respectively). The F1 score is plotted on the *y-axis* and the electrode ID number of the "center" electrode for each 3x3 grid is shown on the *x-axis*. **(A)** Test. **(B)** Validation. **(C)** Online 1. **(D)** Online 2. **(E)** Online 3. Note: the HGLP, HG-C, and HG-PC plots for the results with model trained with permuted labels overlap one another as horizontal lines at approximately 0.4.

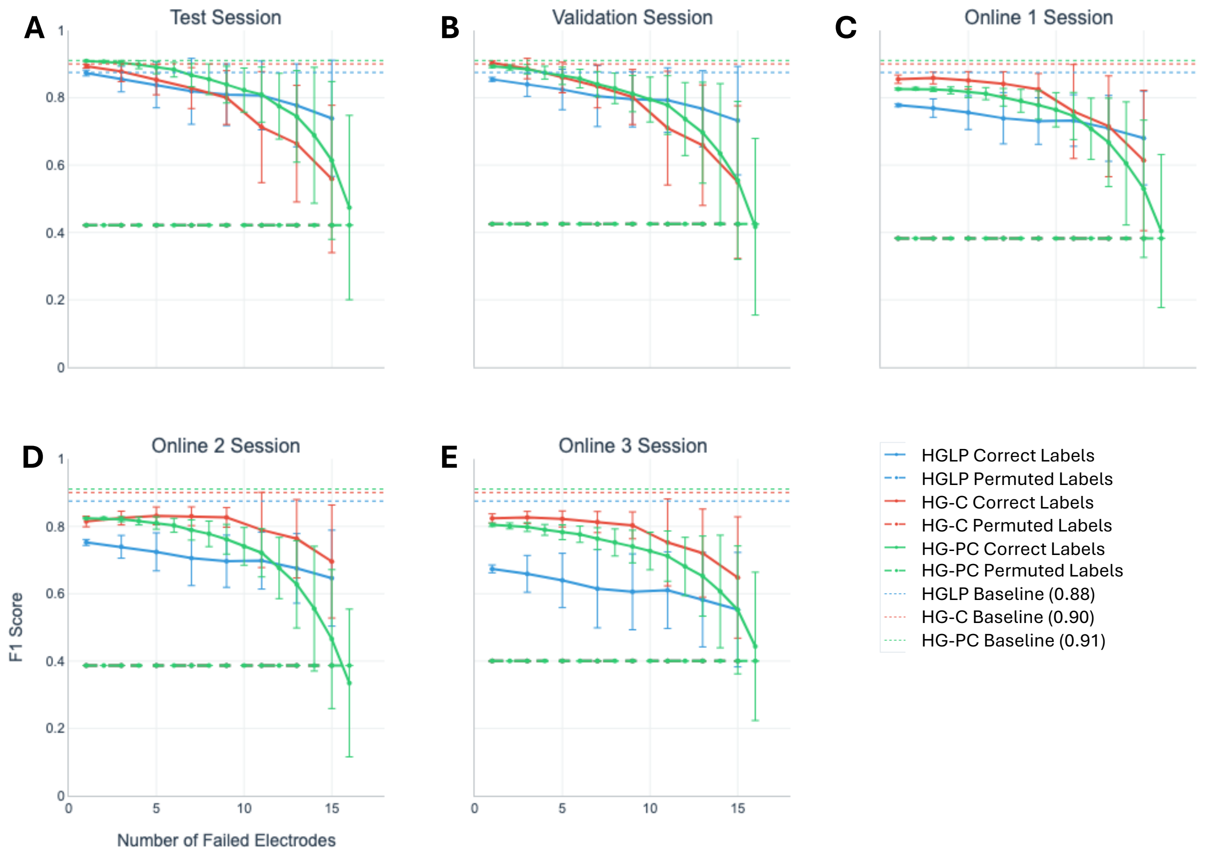

**Figure S9.** Details of the effect of simulated neural signal loss in random subsets of individual electrodes in the ventral grid on NVAD F1 score by session. Neural signal loss is simulated by setting all HGLP values or all pairwise HG-C (HG-PC) values to $N(0, \sigma=1)$ in a random subset of electrodes with subset size ranging from 1 to 16 in steps of 2 and evaluating NVAD performance. For each panel in the Figure – one panel per session - we show: HGLP NVAD trained with correct labels (blue solid line) and an HGLP NVAD model trained with permuted target labels (blue dashed horizontal line at ~0.4); HG-C NVAD trained with correct labels (red solid line) and an HG-C NVAD model trained with permuted target labels (red dashed horizontal line at ~0.4); HG-PC NVAD trained with correct labels (green solid line) and an HG-PC NVAD model trained with permuted target labels (green dashed horizontal line at ~0.4); and the F1 score obtained on Test set by the HGLP, HG-C, and HG-PC NVAD intact models (shown as dotted blue, red, and green lines near the top of each subplot, respectively). The average and standard deviation F1 score is aggregated over repeated runs with different random subsets of the given size and plotted on the *y-axis*. The count of "lost" electrodes is plotted on the *x-axis*. **(A)** Test. **(B)** Validation. **(C)** Online

1. **(D)** Online 2. **(E)** Online 3. Note: the HGLP, HG-C, and HG-PC plots for the results with model trained with permuted labels overlap one another as horizontal lines at approximately 0.4.

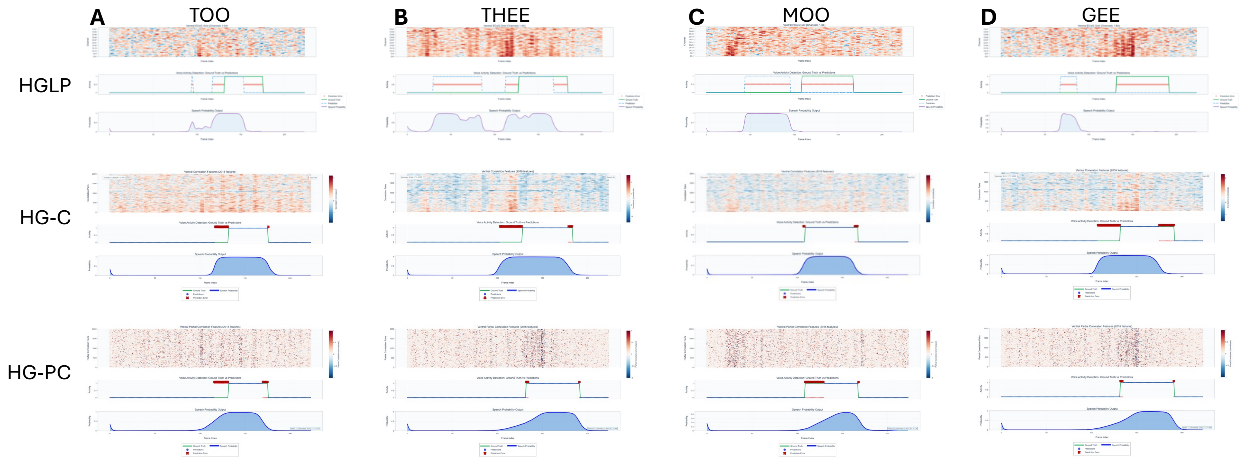

**Figure S10.** Comparison of representative error modes differentiating HGLP, HG-C, and HG-PC performance in the online session 3 approximately 25 weeks following the last training session. **(Column A)** Consonant-vowel pair, “too”. **(Column B)** “Thee”. **(Column C)** “Moo”. **(Column D)** “Gee”. In each column there is one panel each for HGLP (*top*), HG-C (*middle*), and HG-PC (*bottom*). In each panel there are three plots: the feature space representation (*top*); overlay of ground-truth label, predicted label, and indication of errors as orange or red stars (*middle*); and probability of speech frame generated by the NVAD model (*bottom*). All figures are time-aligned (*x-axis*) for ease of reference.
